# Systematic mapping of enhancers, silencers, and topological elements with prime editing deletion screens

**DOI:** 10.64898/2026.10.08.756856

**Authors:** Xiangmeng Shawn Cai, Michael T. Montgomery, Masahiro Nagano, Ashna Nisal, Zhengxi Gu, Ziwei Chen, Kamal Obbad, Ruchir Rastogi, Yingxuan Tan, Tony Zeng, Maya U. Sheth, Coco Xia, Shannon Chiang, Anshul Kundaje, Anders S. Hansen, Lacramioara Bintu, Jesse M. Engreitz

## Abstract

Noncoding regulatory elements in the human genome can control gene expression through distinct mechanisms, including acting as enhancers, silencers, or topological elements. However, we lack tools to identify all such classes systematically and quantify their effects on gene expression in an endogenous genomic context. Here we develop Swap-seq, a method that combines pooled twin prime editing with flow-sorting to delete hundreds of endogenous regulatory sequences and measure their quantitative effects on the expression of a nearby gene. We apply Swap-seq to dissect the regulatory elements that control *PPIF* expression in THP-1 monocytes. We quantify the effects of distal and intronic enhancers, characterize CTCF insulators, and discover a silencer that regulates distal genes through histone deacetylase (HDAC)-mediated repression of neighboring enhancers. Hundreds of other genomic elements share its chromatin signatures and transcription factor binding sites, suggesting that this silencer represents a broader class of repressive elements. We leverage Swap-seq data to benchmark sequence-to-function models and show that they often capture the local activity of regulatory elements but fail to link them to distal target genes. Swap-seq thus provides a generalizable tool to discover regulatory elements across mechanistic classes, characterize their effects on gene expression, and generate large-scale datasets for evaluating sequence-to-function models.

## Introduction

The human genome contains millions of noncoding regulatory elements that control gene expression in development and disease^1,2^. Identifying these sequences and characterizing their functions is a fundamental challenge in modern biology^3,4^. Several classes of regulatory elements have been defined with distinct functions, including promoters, which regulate transcription initiation^5,6^; enhancers, which recruit transcription factors to activate gene expression^7,8^; CTCF binding sites, which block loop extrusion and mark the boundaries of topologically associating domains^9,10^; and silencers, which repress gene expression but remain among the least studied classes^4,11–13^. One obstacle to studying these regulatory elements is the lack of methods that can identify them, map them to the genes they regulate^14,15^, and measure their endogenous effects across all classes, regardless of the underlying mechanism^16,17^. It also remains unclear to what extent sequence-to-function models^18–21^ can predict the effects of classes of regulatory elements beyond enhancers.

Existing high-throughput technologies have characterized thousands of candidate regulatory elements but have been largely restricted to enhancers and carry important limitations. Reporter assays such as massively parallel reporter assays (MPRAs)^22–24^ and STARR-seq^25,26^ have nominated millions of candidate elements^1^, but they do not determine which genes those elements regulate, nor do they measure endogenous effects on gene expression^14,27–29^. CRISPR interference (CRISPRi) coupled with single-cell or sorting-based readouts can identify distal enhancers and map enhancer–gene connections^25,30–33^, but may be limited in its ability to discover other classes of regulatory elements that are insensitive to its KRAB-mediated epigenetic silencing mechanism^17,34,35^. In theory, a method based on genetic deletions could overcome these limitations and probe any class of regulatory elements^14^. In practice, existing CRISPR–Cas9 screens based on two-guide cutting cannot verify the outcome of editing^34,36–38^ and can introduce unintended indels or inversions, which complicates interpretation^14,37,39^.

Here we developed Swap-seq, a high-throughput pooled method based on twin prime editing^40^, to systematically identify any class of regulatory elements and measure their quantitative effects on the expression of a target gene. Swap-seq deletes candidate elements, replaces them with unique barcode cassettes, and uses fluorescence-based cell sorting followed by barcode sequencing to quantify the effects of each deletion on gene expression^32,41,42^. Through this approach, Swap-seq addresses three challenges that previously limited sensitive measurement of the effects of regulatory elements: (i) Swap-seq uses twin prime editing to generate precise deletions without double-strand DNA breaks, thus significantly lowering the frequency of unintended edits that could affect gene expression^40,43^. (ii) It quantifies the effects of edited alleles directly by sequencing the barcode inserted at each edit site, rather than inferring editing efficiency from guide frequency alone, as is common in most pooled CRISPR and prime editing screens^36,44^. (iii) It utilizes flow-sorting to assess tens of millions of single cells per experiment, providing the statistical power to detect small effect sizes^41^.

We applied Swap-seq to systematically characterize the regulatory elements controlling *PPIF*, a key regulator of mitochondrial permeability. In total, we quantified the effects of 208 edits of 50–250 bp across 24 candidate elements with high reproducibility. Comparing Swap-seq with CRISPRi at the same gene, Swap-seq identified regulatory elements that CRISPRi missed, including enhancers within the gene body and CTCF insulators. Most strikingly, Swap-seq identified a silencer that represses *PPIF* by affecting nearby enhancers, a mechanism that appears to reflect a larger class of elements across the genome. Using Swap-seq measurements, we benchmarked and identified limitations of current sequence-to-function models at connecting distal regulatory elements to their target genes. Swap-seq overcomes key technical barriers to characterize regulatory elements at scale, advancing our understanding of gene regulation and previously overlooked classes of regulatory elements.

## Results

### Swap-seq measures effect sizes for hundreds of deletions with high reproducibility

We developed Swap-seq to introduce hundreds of precise genomic deletions and quantitatively measure each of their effects on the expression of a nearby gene in a pooled manner. To generate precise deletions, Swap-seq uses twin prime editing to delete 50- to 250-bp genomic regions and swap in 71-bp cassettes containing deletion-specific barcodes (**Fig. 1A**). To measure effects on expression, we sort cells into bins of expression using RNA FlowFISH, PCR amplify and sequence the deletion-specific barcodes, then use the frequencies of barcodes across bins to estimate the effect of each deletion on gene expression (**Fig. 1A**).

**Figure 1.**
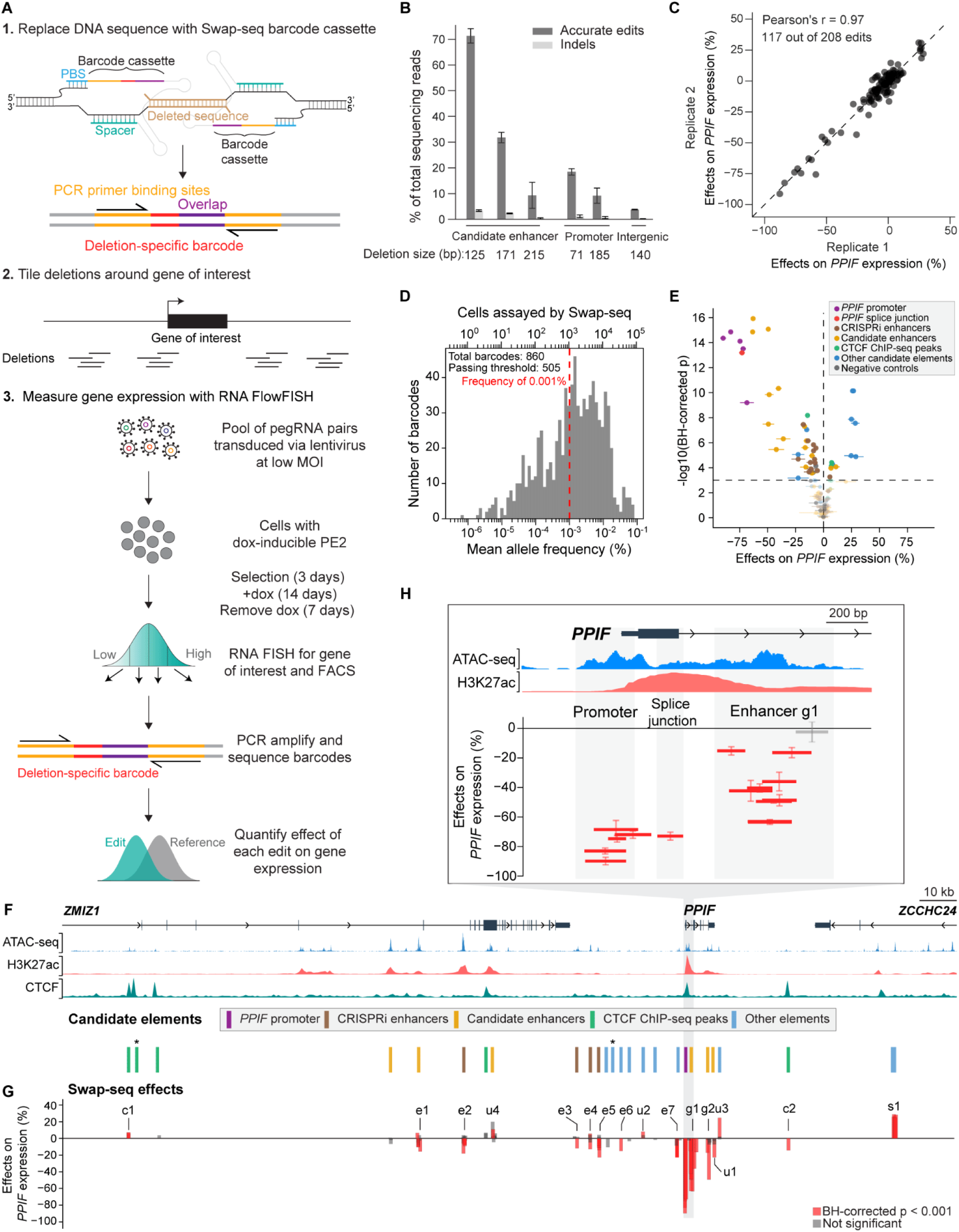
Swap-seq characterizes gene regulatory elements in the *PPIF* locus. **A.** Schematic of the Swap-seq workflow for measuring the effects of DNA sequence deletions on gene expression. Each Swap-seq edit uses a pair of pegRNAs to delete a 50- to 250-bp target sequence and replace it with a barcode cassette containing an edit-specific 8-bp barcode together with sequences common to all edits, including PCR primer-binding sites (yellow) and a 3′ overlap sequence (purple). In a Swap-seq screen, hundreds of such edits are designed to perturb sequences surrounding a gene of interest. Cells are infected with a lentiviral pegRNA-pair library at low multiplicity of infection (MOI); after 14 days of editing, cells are cultured without doxycycline for 7 days before undergoing FlowFISH sorting and quantification of changes in gene expression. **B.** Individual Swap-seq editing test in three regions: the *PPIF* promoter, a candidate enhancer with high accessibility and H3K27ac signal (enhancer e2 in **G**), and a region with low accessibility (hg19: chr10:81,069,000-81,069,517). Error bars: 95% CI for the mean. **C.** Correlation of effect sizes between two biological replicates, after removing edits that did not pass the 0.001% frequency threshold. Effects for each edit are averaged among barcodes. (Pearson’s r = 0.97, n = 117 edits). **D.** Editing efficiency and number of cells assayed for each barcode. A barcode passed the threshold in a given technical replicate if aligned reads exceeded 0.001% of all reads. Bottom axis, mean allele frequency calculated as the percent of reads aligned to the barcode averaged across replicates; y-axis, number of barcodes; top axis, number of cells assayed. **E.** Volcano plot of Swap-seq edits passing the frequency threshold. Edits were considered significant at BH-corrected p < 0.001 (horizontal dashed line). Error bars: 95% CI for the mean. **F.** Swap-seq screen candidate elements at the *PPIF* locus. In total, 24 candidate regulatory elements were targeted and the *PPIF* promoter was included as a positive control. *Two elements with no edits above the frequency threshold of 0.001%. **G.** Swap-seq effects on *PPIF* expression. Red, significant edits (BH-corrected p < 0.001, one-sample t-test); gray, non-significant edits. **H.** Swap-seq edits spanning the promoter, splice junction, and enhancer g1. Red, significant edits (BH-corrected p < 0.001, one-sample t-test); gray, non-significant edits.

We chose the *PPIF* gene locus as our model for applying the Swap-seq workflow. *PPIF* is a non-essential gene that is located in a risk locus for autoimmune and inflammatory diseases^45^ and quantitatively tunes mitochondrial function in THP-1 monocytes^46^. *PPIF* is predicted to be among the top 10% of genes with the most complex enhancer landscapes across cell types^45^, and we have previously identified 5 regulatory elements using CRISPRi-FlowFISH^45^. Thus, the *PPIF* locus provides an ideal test case to determine whether Swap-seq can identify known regulatory elements and discover new ones. To prototype Swap-seq, we first designed a barcode cassette that can be efficiently inserted with twin prime editing (**fig. S1A**). We optimized reverse transcription (RT) templates with different lengths and PCR primer binding sites to identify cassettes with good efficiency and fidelity (**fig. S1B-D**). We then used ChromBPNet^47^, a model that predicts chromatin accessibility from DNA sequence, to design cassettes that are not expected to contain transcription factor binding sites or affect chromatin accessibility in THP-1 cells (**fig. S2**). We tested Swap-seq editing efficiency at three candidate regulatory elements at the *PPIF* locus and observed editing efficiencies ranging from 3.6-71.5% for the intended deletions (71-215 bp) with low indel rates (**Fig. 1B**).

We then designed a Swap-seq paired-pegRNA library to target the *PPIF* promoter, first splice junction, and 24 candidate regulatory elements, including (i) four accessible elements that we previously classified as distal enhancers using CRISPRi-FlowFISH^32,41,45^, (ii) six additional candidate enhancers that are accessible and marked by strong H3K27ac signal, including three elements in the *PPIF* gene body, (iii) five elements observed to bind CTCF by ChIP-seq^48^, and (iv) nine other accessible elements (**Fig. 1F**). We also included the promoters of the two nearby genes, *ZMIZ1* and *ZCCHC24*, to assess potential trans-effects, as several *PPIF* candidate regulatory elements are within their gene bodies. Finally, we included two types of negative controls: (i) edits in accessible regions not on the same chromosome as *PPIF* (“out-of-chromosome” controls), and (ii) non-editing controls, which are constructs that do not contain pegRNAs and instead directly encode the Swap-seq cassette **(fig. S3A**). In total, we designed 208 deletion edits (50–250 bp), each with at least four independent barcode cassettes, to produce a library of 860 pegRNA pairs (**table S2**).

We cloned these pegRNA pairs into a plasmid library and transduced it into THP-1 cells engineered to express the PE2 CRISPR prime editor under the control of a doxycycline-inducible promoter^41^. We induced prime editing for 14 days, then waited an additional 7 days for PE2 degradation to avoid potentially confounding effects of its binding at target loci (**Fig. 1A**). We sorted these cells into four bins of *PPIF* expression using RNA FlowFISH^32,45^. Then, for each bin we extracted genomic DNA, PCR amplified and sequenced the barcode cassettes, and calculated the effect of each barcode-deletion combination using maximum likelihood estimation across the bins while taking into account the total editing efficiency (**Methods**).

We obtained quantitative measurements for 117 out of 208 edits (**table S3**), which were highly reproducible between biological (Pearson’s r = 0.97, **Fig. 1C**) and technical replicates (Pearson’s r = 0.90-0.97, **fig. S4A**) and across the four barcodes for each edit (Pearson’s r = 0.95, **fig. S2G**). Editing efficiency varied across pegRNA constructs, with 505 of 860 barcodes detected at an allele frequency above 0.001% — a threshold corresponding to roughly 1,000 assayed cells per barcode (**Fig. 1D, fig. S4B**). Edits at the *PPIF* promoter reduced expression by 69–90% (**Fig. 1F-H)**, whereas edits across the *ZMIZ1* and *ZCCHC24* promoters did not **(fig. S5**). None of the barcode-only controls affected *PPIF* expression, confirming that measured effects arise from the deletions rather than from construct delivery or readout bias (**fig. S3B**). We generated clonal cell lines carrying homozygous Swap-seq deletions at the enhancer e2 (**Fig. 1G**) and promoter of *PPIF*: qPCR of these clones matched the pooled measurements (**fig. S4C**), confirming that Swap-seq reports quantitatively accurate effect sizes.

Across the locus, Swap-seq discovered 48 edits in the *PPIF* promoter, the splice junction, and 16 other regulatory elements that significantly affected *PPIF* expression (BH-corrected p < 0.001) (**Fig. 1E, table S3**). Four elements had classical enhancer signatures: they were accessible, marked by H3K27ac, not bound by CTCF, and their deletion decreased *PPIF* expression. Of these candidate enhancers, two were distal (e1 and e2), and the other two were in the *PPIF* introns (g1 and g2, **Fig. 1G**). Five other elements that were accessible but not strongly marked by H3K27ac or CTCF (e3-e7, **Fig. 1G**) also led to a decrease in expression upon deletion. Two CTCF-bound elements lacking H3K27ac (c1 and c2) had positive and negative effects on *PPIF* expression, respectively (**Fig. 1G**). All deletions in one element located in an intron of *ZCCHC24* led to significant upregulation of *PPIF*, identifying it as a candidate silencer (s1, **Fig. 1G**).

### Swap-seq identifies regulatory elements missed by CRISPRi

To assess the ability of Swap-seq to identify new regulatory elements, we compared the elements targeted by both Swap-seq and previous CRISPRi experiments^45^ (**Fig. 2A**). Swap-seq identified all four distal enhancers discovered by CRISPRi (e2-e5, **Fig. 1G**), plus another seven elements where CRISPRi did not detect significant effects and three within the *PPIF* gene body (**Fig. 2A**). The effect sizes were correlated between the two methods (Pearson’s r = 0.75; **Fig. 2B**), and the Swap-seq edits with the strongest effects overlapped predicted binding sites for transcription factors^47,49^(e.g., e2 and e5 in **Fig. 2C**).

**Figure 2.**
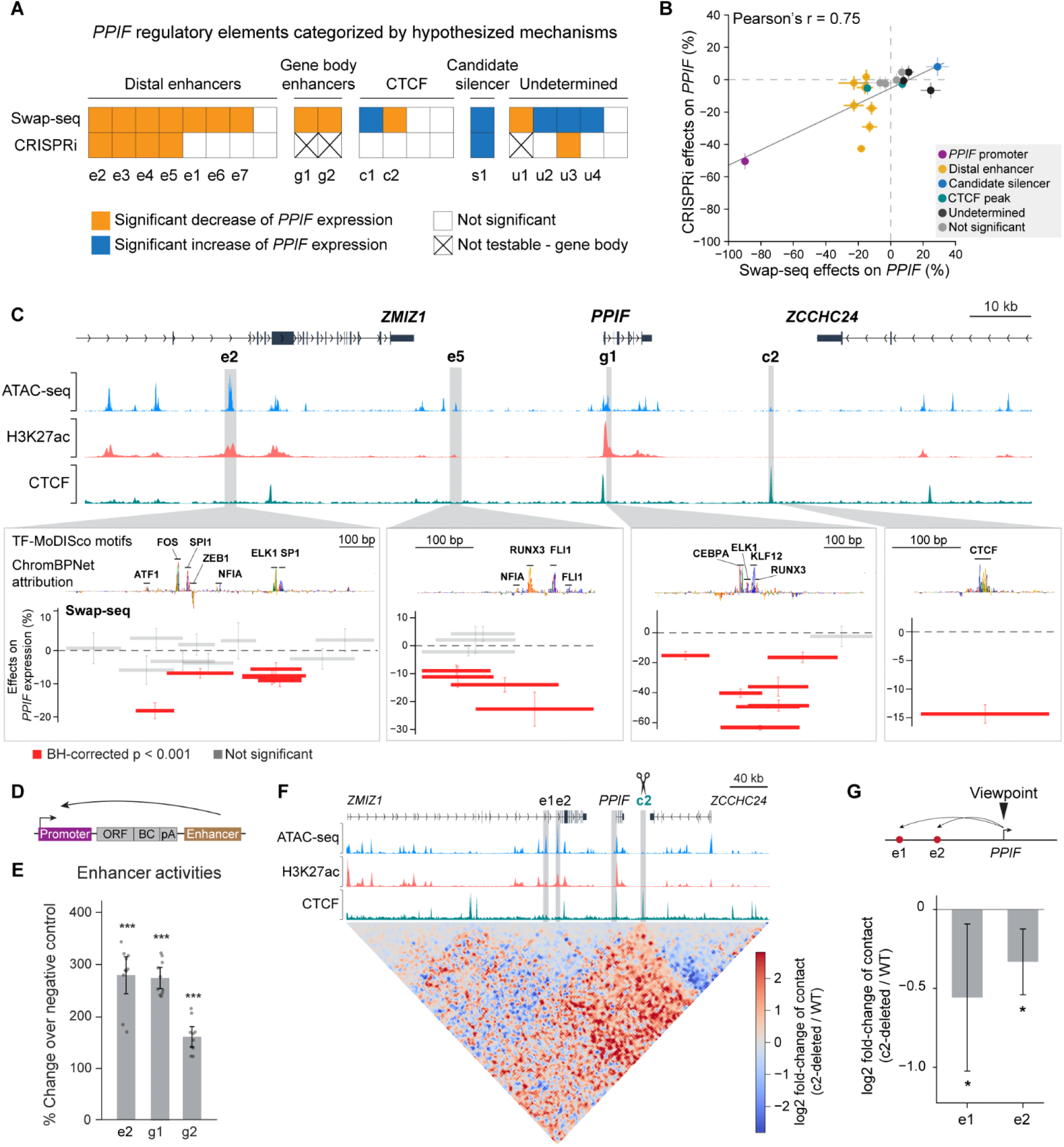
Swap-seq identifies enhancers within the gene body and CTCF insulators. **A.** Comparison of regulatory elements identified by Swap-seq and CRISPRi. An element was classified as significant in Swap-seq if at least one edit within the element was significant (one-sample t-test, BH-corrected p < 0.001). CRISPRi used the average effects of all gRNAs targeting the same region (one-sample t-test, BH-corrected p < 0.05). Elements were grouped according to the hypothesized mechanisms based on their candidate element categories and effects on *PPIF* gene expression. **B.** Correlation between Swap-seq and CRISPRi effect sizes. Elements within the *PPIF* gene body, which are not testable by CRISPRi, were excluded from this analysis; when an element contains several significant edits, the one with the strongest effect was used for comparison with CRISPRi (Pearson’s r = 0.75, n = 18 elements). Error bars: 95% CI for the mean. **C.** Swap-seq edits overlap key transcription factor binding sites (TFBS) within regulatory elements. Close-up views of four elements with significant effects on *PPIF* expression (left to right: e2, a *PPIF* enhancer with high H3K27ac; e5, a low-H3K27ac enhancer; g1, an enhancer in intron 1 of *PPIF*; c2, a CTCF binding site). ChromBPNet attribution scores are plotted together with TF-MoDISco motif annotations. All edits that passed the frequency threshold are plotted with width corresponding to the deleted region. Significance determined by one-sample t-test: red, significant edits (BH-corrected p < 0.001); gray, non-significant edits. Error bars: 95% CI for the mean. **D.** Schematic of the downstream lentiMPRA. Candidate regulatory elements were cloned downstream of the promoter and an ORF–barcode–poly(A) cassette (ORF-BC-pA). Cells were infected with the lentiviral library at high MOI, and barcodes were sequenced from both DNA and RNA; CRE activity was calculated as the ratio of RNA to DNA barcode abundance. **E.** LentiMPRA validation of candidate enhancers. Each dot represents a barcode from two biological replicates. A sequence lacking predicted TFBS (noTFBS) served as a negative control for basal promoter strength. Significance determined by two-sample t-test: *p < 0.05, **p < 0.01, ***p < 0.001. Error bars: 95% CI for the mean. **F.** CTCF deletion alters 3D genome contacts at the *PPIF* locus. Contact frequencies were averaged across region-capture Micro-C (RCMC) measurements from three independent clones, and log2 fold-change was calculated as log2(c2-deleted clones / WT clones) at 1.6 kb bin sizes. **G.** Quantification of contact changes associated with CTCF deletion, with the viewpoint on the *PPIF* promoter. Bars show changes in contact frequency between the *PPIF* promoter and each regulatory element, averaged across three clonal lines at 1.6 kb binsize. Significance determined by two-sample t-test: *p < 0.05, **p < 0.01, ***p < 0.001. Error bars: 95% CI for the mean.

Swap-seq identified three types of candidate elements:

1. Elements that had negative effects on *PPIF* expression when deleted, including three new distal enhancers not detected in previous CRISPRi screens (e1, e6, e7) that contain ChromBPNet-predicted binding sites for activating transcription factors (**fig. S6A**). Additionally, Swap-seq identified two enhancers in the gene body of *PPIF* (g1 and g2, **Fig. 2C**, **fig. S6E-F**), which cannot be identified by CRISPRi as it generally suppresses transcription when targeted to the gene body^30^. To validate these intronic enhancers, we tested g1 and g2 in an enhancer reporter assay^50^ (**Fig. 2D, fig. S7A-C**). Both g1 and g2 strongly activated the *PPIF* promoter in the reporter assay, supporting their functions as enhancers (**Fig. 2E, fig. S7H, table S4**). Another element within the 3’ UTR of *PPIF* (u1) showed decreased expression when deleted, but we categorized it as “undetermined” because it is possible that this UTR deletion may affect *PPIF* at the post-transcriptional level (**fig. S6G**).
2. Two new elements were CTCF binding sites (c1 and c2), where CRISPRi may not have been able to displace CTCF binding^17,51^(**Fig. 2C, fig. S8A-B**). We generated clonal cell lines carrying homozygous deletions of the c2 CTCF site, and conducted region capture Micro-C (RCMC)^52^ to understand its mechanism (**Fig. 2F, fig. S8C**). c2 is downstream of the *PPIF* gene at the boundary of the contact domain containing the gene. Deleting c2 led to a 14.4% decrease in *PPIF* expression (**Fig. 2C, right**) alongside a significant loss of insulation (**Fig. 2F, fig. S8C-D**). Consistent with the observed decrease in *PPIF* expression, contact between the *PPIF* promoter and distal enhancers (e1 and e2) decreased upon c2 deletion (**Fig. 2F-G**). Additionally, we observed increased contact between the candidate repressive element s1 and the *PPIF* promoter and distal enhancers (e1, e5), which can potentially lead to decreased enhancer activities (**Fig. 2F, fig. S8E**).
3. Four elements had positive effects on *PPIF* upon deletion, which we assigned as candidate silencers (**Fig. 2A-B**). We classified three of these elements as “undetermined,” because they contained only one significant edit each (u2, u3, and u4, **fig. S6B-D**), with CRISPRi and Swap-seq showing discordant directions of effect at one (u3, **Fig. 2A**), although notably some did overlap with predicted binding sites for transcription factors with known repressive function^53–55^(**fig. S6B-D**). We focused further on candidate silencer s1, where all 5 tiled deletions led to an increase of *PPIF* expression by 25-29% (**Fig. 3A**).

**Figure 3.**
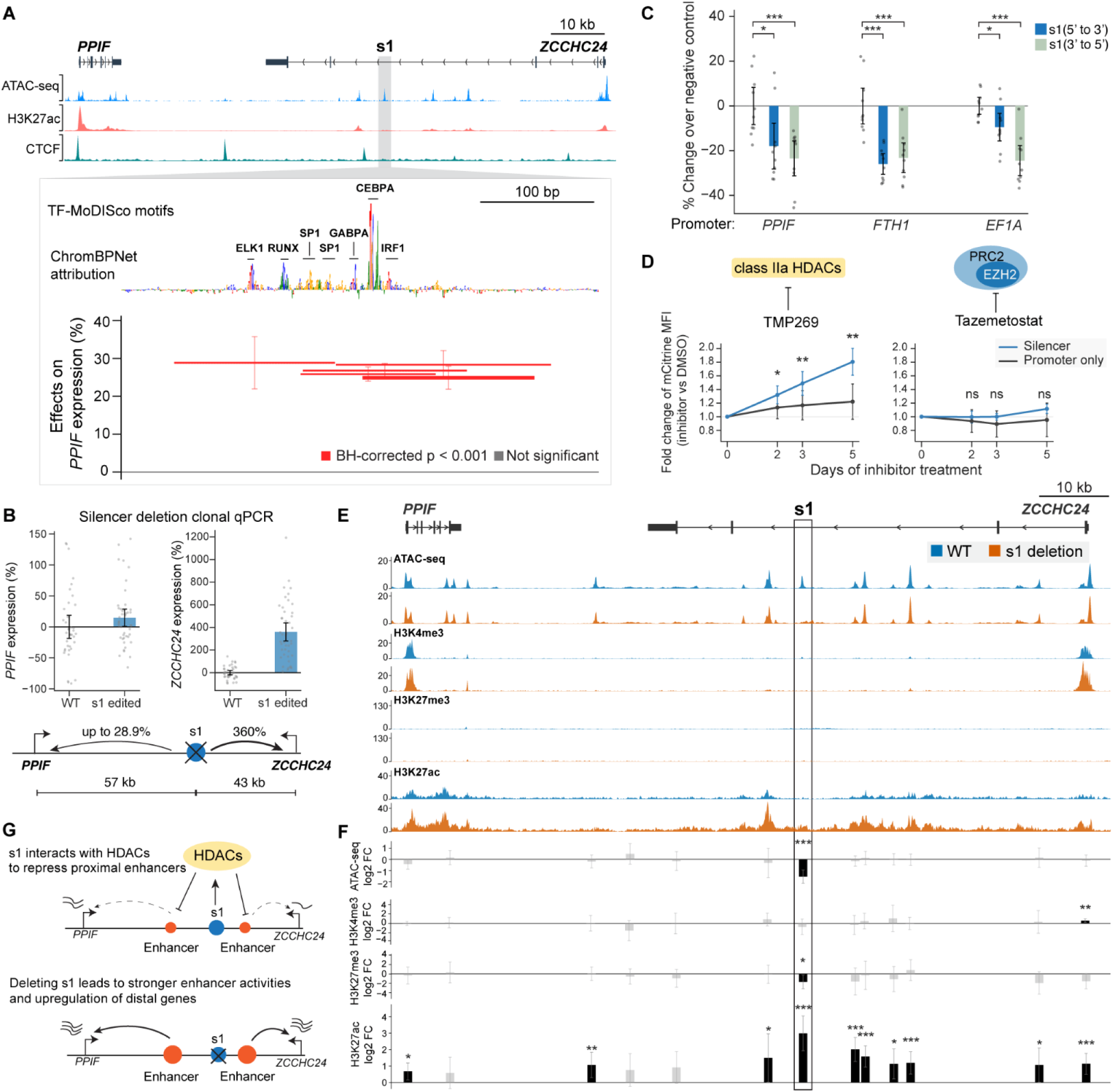
Swap-seq identifies a silencer that represses enhancers through histone deacetylation. **A.** Locus plot of the silencer s1, located within an intron of *ZCCHC24* (hg19: chr10:81,104,100-81,207,000). Close-up shows the ChromBPNet attribution track, annotated TFBS, and Swap-seq edits within s1. Significance determined by one-sample t-test: red, BH-corrected p < 0.001; gray, not significant. Error bars: 95% CI for the mean. **B.** qPCR quantification of the effects of s1 edits on *PPIF* and *ZCCHC24* gene expression. qPCR was used to measure the change of *PPIF* and *ZCCHC24* gene expression between WT clones and clones carrying a homozygous Swap-seq edit of s1. Each dot represents results from an independent clone, averaged between three qPCR replicates. Error bars: 95% CI for the mean. **C.** Validation of the silencer by lentiMPRA. The silencer, in both orientations, was paired with three promoters: *PPIF*, *EF1A*, and *FTH1*. A sequence lacking predicted TFBS (noTFBS) served as a negative control for basal promoter strength. Significance determined by two-sample t-test: *p < 0.05, **p < 0.01, ***p < 0.001. Error bars: 95% CI for the mean. **D.** Inhibitor treatment of cells with lentiviral fluorescent reporter construct. Cells were treated with the indicated inhibitors over a 5-day time course and assayed on days 0, 2, 3 and 5. Dots show mean fluorescence intensity (MFI), expressed as fold-change over DMSO controls, averaged across 3 biological replicates. Blue, reporter construct with silencer; gray, *EF1A* promoter-only control. Significance determined by unpaired two-sided Welch’s t-test: *p < 0.05, **p < 0.01, ns, not significant. Error bars: 95% CI for the mean. **E.** Genomic tracks of accessibility and histone modifications at the silencer s1 locus in WT and clonal lines carrying s1 deletion. Blue, WT; orange, homozygous edited clones. Top to bottom: ATAC-seq (WT, 4 clones; Edited, 6 clones), H3K4me3 CUT&RUN (WT, 3 clones; Edited, 3 clones), H3K27me3 CUT&RUN (WT, 6 clones; Edited, 6 clones), and H3K27ac CUT&RUN (WT, 3 clones; Edited, 3 clones). The scale of H3K27me3 is set such that the maximum matches signal at the exemplary locus enriched with this signal (see **fig. S9C**). **F.** Quantification of ATAC-seq and CUT&RUN signal changes between WT and clones with the s1 edit. Significance determined by Wald z-test: *p < 0.05, **p < 0.01, ***p < 0.001, ns, not significant. Peaks with significant changes highlighted in black. Error bars: 95% CI for the mean. **G.** Model of silencer s1 mechanism. The silencer depends on HDACs to repress nearby enhancers that regulate *PPIF* and *ZCCHC24*. Deletion of the silencer leads to increased enhancer activities, thereby upregulating *PPIF* and *ZCCHC24*.

### A distal silencer represses *PPIF* and nearby enhancers

The candidate silencer element s1 corresponded to an accessible peak in the intron of *ZCCHC24*, 57 kb from the *PPIF* promoter. All deletions within s1 led to an increase in *PPIF* expression (25%-29%; **Fig. 3A**). This element is particularly interesting because the magnitude of its effects was comparable to *PPIF* enhancers, but in the opposite direction; because its chromatin signature did not match any of those previously reported to mark transcriptional silencers^16,56–59^; and because silencers have not been reported for genes as highly expressed as *PPIF*. Accordingly, we sought to determine whether s1 indeed acted as a *cis-*acting silencer, and if so by what mechanism.

We first sought to determine whether the effect on *PPIF* expression was indeed a direct effect, rather than an indirect effect of the element regulating its overlapping gene, *ZCCHC24*, whose RNA or protein products might then affect *PPIF* expression in *trans* (**fig. S9A, Model 1**). To examine possible effects on *ZCCHC24* expression, we generated clonal cell lines homozygous for the Swap-seq edit of s1 and performed qPCR for *PPIF* and *ZCCHC24*. This confirmed our previous observation of an increase in *PPIF*, and revealed a large, 360% increase in *ZCCHC24* expression (**Fig. 3B**). However, downregulation of the *ZCCHC24* promoter via Swap-seq deletions or CRISPRi also led to an increase of *PPIF* expression (**fig. S5**). These contradictory directions of effects between *PPIF* and *ZCCHC24* (specifically, downregulation of *ZCCHC24* via promoter perturbation and upregulation via s1 deletion both increase *PPIF*) are inconsistent with *ZCCHC24* gene products mediating the observed effects of s1 on *PPIF*, suggesting a true *cis*-acting mechanism.

We next considered previously reported mechanisms by which a distal element like s1 might act as a cis-regulatory element to reduce *PPIF* expression: (1) repressing a promoter via direct 3D contact^59,60^; (2) seeding a repressive chromatin domain (Polycomb or heterochromatin)^57,60,61^; or (3) acting as a promoter and thereby reducing expression of a nearby gene via promoter competition^30,62,63^ or transcriptional interference^11,64,65^ (**fig. S9A**, **Models 2-4**, respectively).

Upon examining CUT&RUN, ChIP-seq^66^, RNA-seq^66^, and CAGE^67^ data in THP-1, we found that none of these models was consistent with the chromatin state of the locus. Element s1 did not exhibit strong focal contacts with the promoter of *PPIF* (**fig. S9D**), nor was it strongly marked by either H3K27me3 or H3K9me3 (**fig. S9B-C**). It also did not show substantial evidence of transcriptional activity, based on H3K4me3 and H3K27ac CUT&RUN, RNA-seq, and CAGE (**fig. S9B**). These results led us to consider an alternative model, where s1 can regulate distal gene expression by repressing nearby enhancers (**fig. S9A, Model 5**).

To determine whether s1 is sufficient to repress gene expression, we cloned s1 into a lentiMPRA^50,68^ construct downstream of the reporter cassette and tested its effect on the promoters of *PPIF*, *EF1A*, and *FTH1* **(fig. S7A-E, table S4)**. Indeed, s1 repressed all three promoters by up to 25%, in both forward and reverse orientations **(Fig. 3C)**. We next cloned s1 upstream of gene promoters in a lentiviral fluorescence reporter construct **(fig. S10A-C)**, and performed 5’-end RNA sequencing **(fig. S10D)**. Again, s1 strongly repressed reporter expression, and we did not detect any reporter transcripts originating from s1 (**fig. S10D**). Together, this evidence demonstrates that s1 has autonomous silencing activity and does not act as a promoter.

To identify potential co-repressors required for the silencer’s effect, we tested s1 in the fluorescence reporter assay (**fig. S10A-C**), with and without the addition of drugs targeting specific repressive complexes: tazemetostat, an inhibitor of EZH2^69^, the core catalytic subunit of Polycomb Repressive Complex 2 (PRC2)^70^; and TMP269^71^, an inhibitor of class IIa HDACs (**Fig. 3D**). Over five days of treatment, cells carrying the silencer construct showed significant derepression upon TMP269 treatment relative to the DMSO control (1.8-fold), whereas cells carrying the promoter-only construct showed no significant change (**Fig. 3D**). In contrast, EZH2 inhibition did not significantly change expression from the silencer construct (**Fig. 3D**). This suggests that s1’s repressive activity depends on class IIa HDACs rather than PRC2, consistent with the lack of strong H3K27me3 signal at this locus (**fig. S9B-C**).

To further investigate how this silencer affects *PPIF* expression in its endogenous context, we conducted CUT&RUN and ATAC-seq in wild-type cells and clones carrying a homozygous Swap-seq edit at s1 (**Fig. 3E**). Deleting s1 led to a significant increase of H3K27ac signal at several accessible peaks surrounding the silencer (within 30 kb on each side, **Fig. 3F**). In contrast, neither accessibility nor H3K27me3 changed significantly outside of the site of deletion (**Fig. 3E-F**).

Thus, s1 has autonomous repressive activity that is orientation-independent, is reversible by class IIa HDAC inhibition, and reduces H3K27ac at nearby elements in the genome. Together, these observations suggest a novel model for the silencing activity of s1: it acts as a silencer for nearby genes not via direct 3D contact with distal promoters, but rather by repressing nearby enhancers through HDAC-dependent deacetylation (**Fig. 3G**).

### Sequence syntax of the distal silencer s1

To identify the TFBS responsible for the silencer’s repressive effects, we conducted tiling mutagenesis with Variant-EFFECTS^41^(**Fig. 4A**). We deleted every 5-bp window across a 200-bp region centered on s1, designed edits to disrupt candidate TFBS predicted by ChromBPNet (**Fig. 4B**), and measured the effect of each edit on *PPIF* expression (**table S5**).

**Figure 4.**
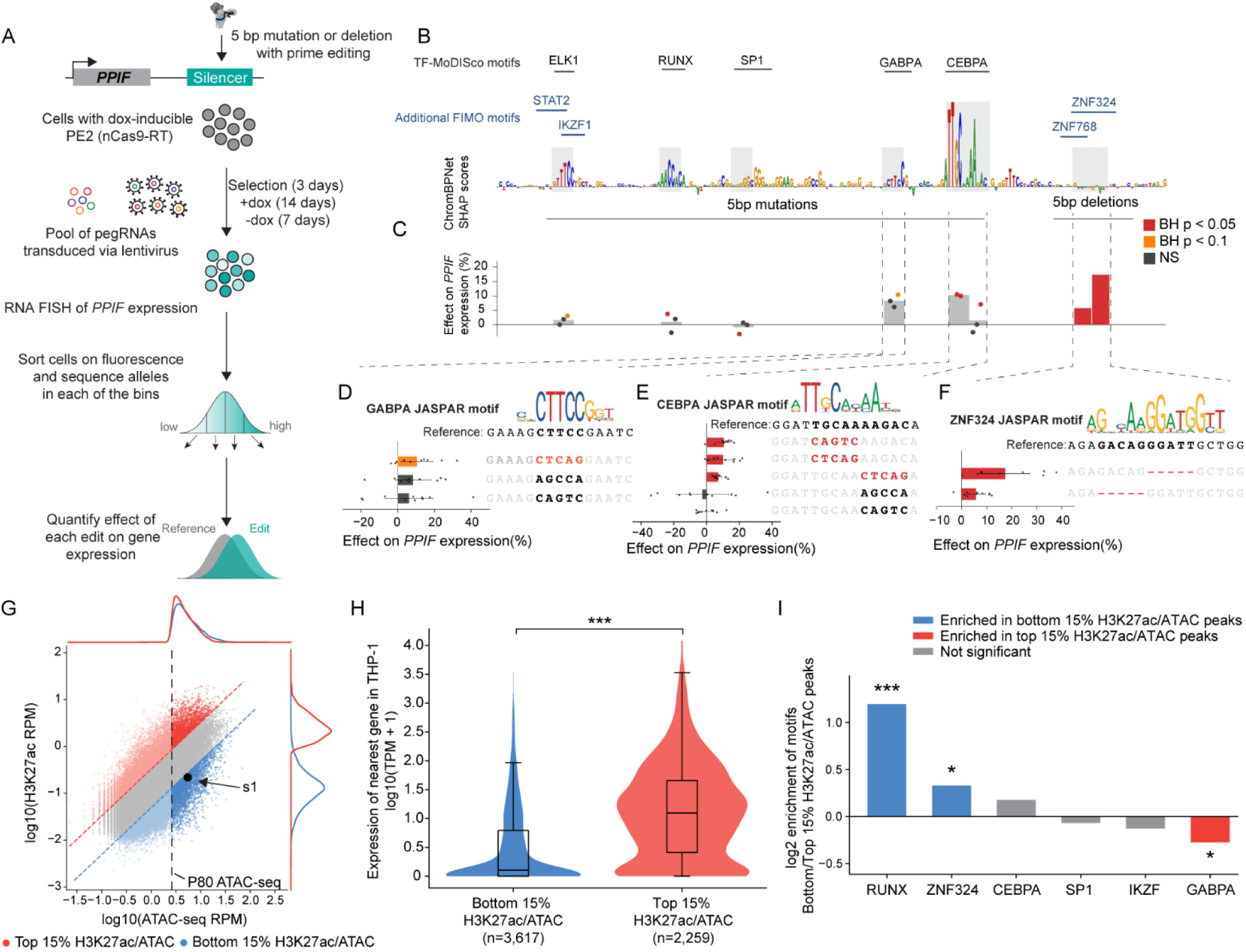
Dissecting the s1 sequence syntax and nominating silencer-like elements genome-wide. **A.** Schematic of the Variant-EFFECTS screen at the *PPIF* silencer. Cells were infected with a lentiviral library of pegRNAs designed to delete or mutate 5-bp sequences across the silencer. Following editing as previously described^41^, gene expression was measured by RNA FlowFISH and sequencing. **B.** ChromBPNet attribution scores and TF-MoDISco annotation of TFBS within the silencer. Blue, additional TFBS annotated by FIMO^78,79^. **C.** Variant-EFFECTS results. For mutations, each bar represents the average effect of three independent 5-bp mutations, shown as dots; for deletions, bars represent two consecutive 5-bp deletions. Significance determined by one-sample t-test: Red, BH-corrected p < 0.05; orange, BH-corrected p < 0.1; gray, not significant. Error bars: 95% CI for the mean. **D.** Effects of 5-bp mutations within the GABPA motif. The reference sequence of this region is shown below the JASPAR position weight matrix (PWM) for the GABPA motif, with the mutations shown beneath. Bar plots show the effect of each mutation; dots represent individual FlowFISH replicates. Significance determined by one-sample t-test: Red, BH-corrected p < 0.05; orange, BH-corrected p < 0.1; gray, not significant. Error bars: 95% CI for the mean. **E.** As in **D**, but for mutations within the CEBPA motif. **F.** Effects of 5-bp deletions matching the ZNF324 motif. The reference sequence is shown below the ZNF324 PWM, with deletions represented as dashed lines. Significance determined by one-sample t-test: Red, BH-corrected p < 0.05; orange, BH-corrected p < 0.1; gray, not significant. Error bars: 95% CI for the mean. **G.** Genome-wide silencer discovery. We excluded peaks below the 80th percentile of ATAC-seq signal and classified peaks into silencer-like or enhancer-like by using bottom and top 15% H3K27ac/ATAC-seq ratio, respectively. Marginal distributions of ATAC-seq and H3K27ac signals are shown on the corresponding axes. Dashed line, ATAC-seq P80. Blue, silencer-like peaks with bottom 15% H3K27ac/ATAC-seq ratio. Red, enhancer-like peaks with top 15% H3K27ac/ATAC-seq. **H.** Comparing expression of genes near silencer-like (bottom 15% H3K27ac/ATAC) and enhancer-like (top 15% H3K27ac/ATAC) peaks. p=2.2x10^-214^, Mann-Whitney U test. **I.** Motif enrichment analysis of TFBS using FIMO and JASPAR and HOCOMOCO databases. Enrichment of motif instances was compared between silencer-like (bottom 15% H3K27ac/ATAC) and enhancer-like (top 15% H3K27ac/ATAC) peaks. Two groups were filtered so they have similar distributions of distances to the nearest gene (see **Methods**). Significance determined by Fisher’s exact test: *BH-corrected p < 0.05, **BH-corrected p < 0.01, ***BH-corrected p < 0.001, ns, not significant.

Edits that disrupted predicted binding sites for RUNX and CEBPA led to significant increases in *PPIF* expression (+3.7% to +10.5%, BH-corrected p = 7.9 × 10^-4^ to 0.027; **Fig. 4C-E**). Notably, ChromBPNet predicted that both of these motifs contributed positively to chromatin accessibility, and both factors have been observed to have either activating or repressive functions in different contexts^72–77^. Additionally, two adjacent 5-bp deletions within a sequence closely matching the canonical ZNF324^74^ motif resulted in an increase in *PPIF* expression of up to 17.4% (**Fig. 4F**). This sequence was not prioritized by ChromBPNet, possibly because the model is known to miss motifs for some TFs^47^. Together, these results suggest that motif instances for RUNX, GABPA, CEBPA and ZNF324 contribute to the ability of the silencer to repress gene expression.

We wondered whether other similar silencers might exist in other loci. We considered the most accessible elements in the genome, and compared two categories: those with low H3K27ac relative to their level of accessibility (similar to the silencer s1), versus those with high H3K27ac (putative enhancers, **Fig. 4G**). The nearest genes of silencer-like peaks (low H3K27ac/ATAC ratio) had substantially lower expression than the nearest genes of enhancer-like peaks (**Fig. 4H**). Motif analysis showed that motifs including RUNX and ZNF324 were significantly enriched in silencer-like peaks (**Fig. 4I**), and 160 peaks in this class contained at least 3 of the 4 motifs (RUNX, GABPA, CEBPA, and ZNF324; **table S6**), suggesting s1 represents a broader class of human silencers.

### Benchmarking sequence-to-function models

The diverse types of regulatory elements identified by Swap-seq, including transcriptional silencers and CTCF sites, provide a unique opportunity to benchmark sequence-to-function models such as AlphaGenome^18^, Enformer^19^, and Borzoi^21^. While these models hold promise for predicting gene expression directly from DNA sequence, evaluation has been challenging due to the lack of large gold-standard datasets linking DNA variants to effects on gene expression^41,80^.

We compared Swap-seq measurements to model predictions across 48 edits in 18 elements that had significant effects (BH-corrected p<0.001) on *PPIF* expression (**fig. S11A, table S7**). Correlation between model prediction and Swap-seq effects was strong (Pearson’s r = 0.64-0.78, **fig. S11B**), but this was driven by strong performance on elements <10 kb from the promoter (**Fig. 5A**). For elements >10 kb from the promoter, model-predicted effects were poorly correlated with experimental effects (Pearson’s r ranging between –0.12 for AlphaGenome THP-1 CAGE and 0.27 for Enformer CD14+ monocyte CAGE, **Fig. 5B-C**, **fig. S11C-E**).

**Figure 5.**
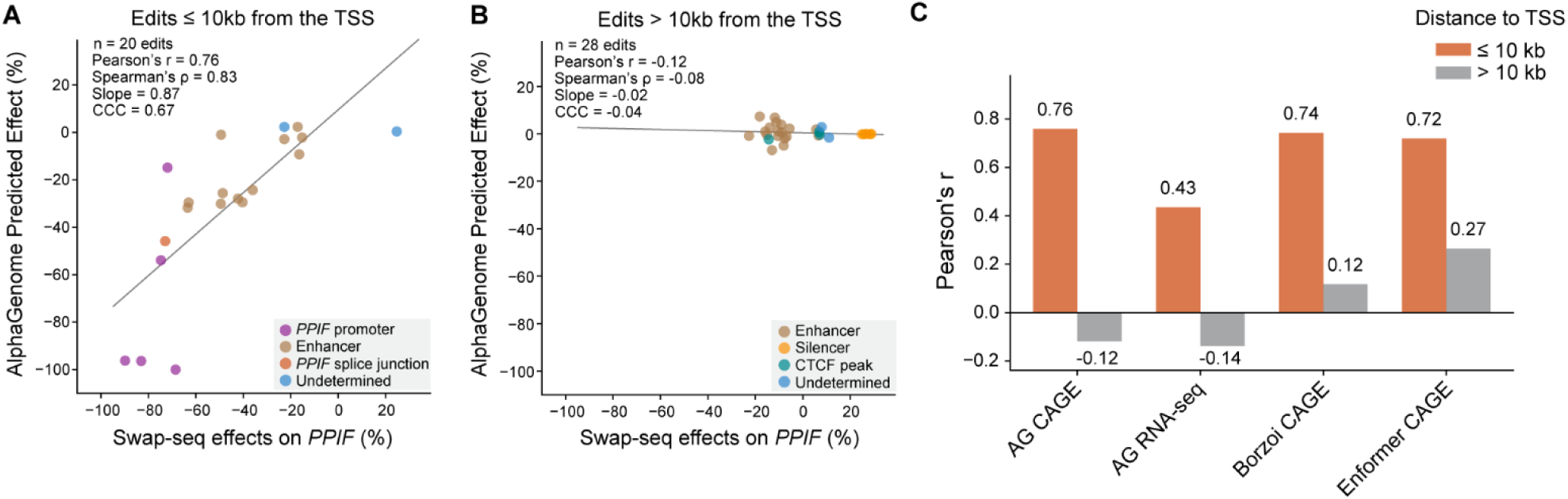
Benchmarking sequence-to-function models using Swap-seq data. **A.** Correlation between Swap-seq effects and AlphaGenome predictions (1-Mb context, THP-1 CAGE track) for edits within 10 kb of the *PPIF* TSS (Pearson’s r = 0.76, n = 20 edits). **B.** As in A, but for edits more than 10 kb from the *PPIF* TSS (Pearson’s r = −0.12, n = 28 edits). **C.** Pearson correlation coefficient between Swap-seq effects and model predictions (AlphaGenome THP-1 CAGE, AlphaGenome CD14+ monocyte RNA-seq, Borzoi CD14+ monocyte CAGE, and Enformer CD14+ monocyte CAGE), stratified by distance to the *PPIF* TSS.

We more closely interrogated the ability of the models to detect the regulatory activities of distal enhancer e2 and silencer s1. While the models did not detect a long-range effect of these regulatory elements in their endogenous locus (**Fig. 5B**), we wondered whether the models might detect their local regulatory activities. To test this, we used AlphaGenome to predict promoter CAGE signals for the same MPRA reporter sequences we measured experimentally (**Fig. 2E**, **Fig. 3C**), with regulatory elements inserted downstream of the transcript at a much shorter distance from the promoter than in the endogenous locus. Although AlphaGenome captured the enhancer activation at this proximal location, its predictions on the silencer did not agree with the MPRA results and were inconsistent: it predicted repressive effects on some promoters, and activating effects on others (and different effects in different sequence contexts) (**fig. S12)**. Thus, AlphaGenome captures local activating effects of enhancers but not the repressive effects of silencer s1, and does not propagate these effects to a distal target gene. Future work could apply Swap-seq at scale to build the larger silencer- and enhancer-effect datasets needed to evaluate and fine-tune such models.

## Discussion

Deletion-based direct mapping of regulatory element-gene interactions has been a long-standing goal in the genomics field^14^. Early two-guide CRISPR–Cas9 approaches, applied individually or in pooled screens, suffered from limited sensitivity and specificity due to unintended editing outcomes — indels, inversions, scars and insertions^39^ — and pooled formats could assess neither editing outcome nor efficiency for any given deletion pair^14^. Recent work used twin prime editing screens to identify large sequence deletions that affected cellular fitness, which improved editing precision over previous CRISPR-Cas9 approaches, but it could not measure the quantitative effects of regulatory sequences nor account for variable editing efficiencies^81^. Swap-seq addresses these challenges by using pooled twin prime editing^40^ with barcode readout to enable tiling deletion screens that can assess the precise quantitative effects of regulatory elements on gene expression, regardless of the mechanism (**Fig. 1**).

Swap-seq also addresses some of the long-standing questions about CRISPRi tiling — the workhorse approach for mapping enhancer-gene regulatory interactions over the last decade^30,31,33,35,45,82,83^. There have been two concerns: (i) the possibility of 1D or 3D spreading of CRISPRi epigenetic silencing leading to false positives^31,84–88^ and (ii) that KRAB may be unable to repress certain classes of elements, generating false negatives^34,51^. Our large-scale deletion analysis indicates that (i) does not appear to be an issue—all four distal enhancers discovered by CRISPRi validate by deletion, consistent with previous results showing that individual CRISPRi-detected enhancers validate by genetic deletion^31^; at least when CRISPRi is applied for a short period of time, e.g., <48 hrs, which limits spreading of silencing^89^ (**Fig. 2A, B**). (ii) CRISPRi does appear to miss some types of regulatory elements (*e.g.*, enhancers in the gene body, CTCF sites) (**Fig. 2A**). CRISPRi targeting silencer s1 showed upregulation of *PPIF* — but with an 8% increase, the effect size is much smaller than that measured by Swap-seq (**Fig. 2B**). It is unclear whether CRISPRi can be used to characterize silencers, because dCas9-KRAB is itself a repressor, so recruiting it to DNA creates a synthetic silencer possibly of a different type and with a different range of action^34,35^. Therefore, these two approaches have complementary strengths: CRISPRi remains more scalable for perturbing enhancers and promoters, owing to its higher and more consistent efficiency, whereas Swap-seq casts a broader net across elements with different regulatory mechanisms, most notably silencers.

Silencers are a class of elements that regulate gene expression in *cis* through repressive cofactors^11,12^. They are one of the understudied classes of elements — in large part because we as a field have not discovered and characterized many examples in the genome, and major open questions remain about their regulatory syntax and required cofactors. Many repressive elements have been proposed to decrease gene expression in reporter assays^1,13,16,90,91^, but most of them have not been validated or characterized in their endogenous genomic loci. Among bona fide silencers, the CD4 silencer is a known example of a gene-proximal regulatory element that interacts with RUNX family TFs and HDACs to repress *CD4* expression in a cell-type-specific manner^73,92^. Other examples have shown that silencers could regulate distal genes through spreading of histone modifications^57,93^, or polycomb-mediated 3D contact^56,60^. In contrast, s1 appears to repress distal gene expression through a distinct mechanism — by interacting with HDACs to decrease the activities of silencer-proximal enhancers (**Fig. 3**). Using chromatin features and TFBS of s1, we nominated hundreds of potential silencers in the genome (**Fig. 4**). These results have three further implications: (1) distal silencers may not necessarily need to establish direct contact with the promoters they regulate; (2) silencers can quantitatively tune gene expression, even for highly expressed genes, instead of acting like an on/off switch; and (3) many more silencers likely exist in the genome that would be missed by searches restricted to elements bearing repressive chromatin signatures (e.g., H3K27me3, H3K9me3)^1,11,16,94^.

Sequence-to-function models^18–21^ hold the potential to help discover regulatory elements^13^ and understand their mechanisms^95–97^, as well as advance our ability to dissect genetic diseases^98,99^ and develop gene therapies^41,100–102^. However, to evaluate and improve these models requires large gold-standard datasets^41,80,103^. Swap-seq pairs precise DNA sequence edits with a quantitative readout of gene expression, which we used to benchmark against existing sequence models that predict transcription-related features (**Fig. 5**). Model predictions correlate with Swap-seq effects for elements within 10 kb of the TSS, but none of the models appear to successfully predict the effects of more distal elements, despite all having a sufficient context window in training (up to 1 Mb for AlphaGenome) (**Fig. 5A-C**). Future work will apply Swap-seq to more loci and cell types to generate sufficiently large datasets that can evaluate and improve sequence-based models.

To support future applications of Swap-seq, we note several technical considerations: (1) While Swap-seq is broadly applicable to study a variety of genes via RNA FlowFISH, each experiment is limited to measuring the effects of candidate regulatory elements for one target gene. However, Swap-seq can be adapted to measure cellular phenotypes, such as cell growth or differentiation, which allows its application to regulatory elements genome-wide in one experiment. (2) Swap-seq editing efficiency, like prime editing as previously reported^104^, is affected by chromatin accessibility and histone modifications (**fig. S4D**). Future work is needed to evaluate how this limitation affects applying Swap-seq to target inaccessible gene regulatory elements or repressive elements in heterochromatin^105^. (3) Wide variability in editing efficiency limits the scale of Swap-seq. It is worth noting that we used inducible-PE2 cell lines for this work, and future work can improve editing efficiency with more advanced prime editing tools^106–108^. Higher efficiency editing would expand the scale of Swap-seq screens further.

In summary, Swap-seq enables deletion-based high-throughput characterization of gene regulatory elements. In this work we focused on distal regulatory elements and quantitative measurements of gene expression, but the Swap-seq design can generalize to study how DNA deletions could affect cellular function through other mechanisms, such as by altering protein-coding sequences, splice sites, or RNA regulatory elements^109,110^. Swap-seq is also compatible with screening on proliferation effects^111^, or on other sortable phenotypes such as cell differentiation^112^. We anticipate that Swap-seq will help discover and characterize gene regulatory elements, resolve their mechanisms of action, and generate datasets needed to benchmark and improve sequence-to-function models.

## Methods

### Tissue culture

We used inducible PE2 THP-1 cells that were previously engineered to express PE2-P2A-BFP upon treating cells with doxycycline^41^. We maintained THP-1 cells between 500K to 1.5M cells per mL in RPMI-1640 (Thermo Fisher Scientific) with 10% heat-inactivated FBS (Thermo Fisher Scientific), 4 mM L-glutamine, and 100 units/mL streptomycin and 100 mg/mL penicillin. We maintained HEK293T cells (ATCC) between 20-90% confluence in DMEM with 1 mM sodium pyruvate, 25 mM glucose (Thermo Fisher Scientific) and 10% heat-inactivated FBS. Both cell lines were cultured at 37°C in 5% CO2.

### Swap-seq pegRNA optimization in HEK293T cells

Swap-seq is based on twinPE^40^, a method that uses paired pegRNAs to delete large DNA sequences. Unlike other “seamless” methods such as PRIME-Del^43^, twinPE requires that both pegRNAs encode a complementary sequence on the RT template. Upon PE-mediated reverse transcription, these complementary sequences will form the 3’ overlapping flaps that anneal to replace the deleted sequences. We harnessed the flexibility of twinPE to encode a Swap-seq barcode cassette on RT templates that contains (1) a constant forward PCR primer binding site, (2) an 8-bp deletion-specific barcode, (3) the overlapping sequence, and (4) the reverse PCR primer binding site (fig. S1A).

We tested how different pegRNAs could affect editing efficiency and accuracy. We cloned pegRNAs that delete 90 and 89 bp at HEK3 and CCR5 respectively, using the same spacers that have been reported to have high editing efficiency. We varied the length (12, 16, 21 bp) and GC% (62, 71, 81%) of the overlapping sequence, as well as the PCR primer binding sites of the barcode cassette (**table S1**). To transfect HEK293T cells, we coated 48-well tissue culture plates with 1:20 dilution of Poly-D-lysine solution (Millipore Sigma #A-003-E) and plated 50,000 HEK293T cells per well on Day 0. The next day, we added 125 ng of plasmids carrying each pegRNA, 250 ng of pCMV-PE2A (Addgene #132775), and 1 uL of the X-tremeGENE HP^TM^ DNA Transfection Reagent (Millipore Sigma #06366236001) according to the manufacturer’s instructions. We harvested the genomic DNA from HEK293T cells 72 hours post-transfection (see **Genomic DNA extraction and barcode amplicon sequencing**). We amplified the edited loci with Q5 High-Fidelity 2X Master Mix (New England Biolabs #M0492L) and 0.5 uM of each amplification primer (**table S1**) and sequenced the amplicon on an Illumina sequencing platform. Sequencing data were analyzed using CRISPResso2^113^ as previously described in Anzalone et al.^40^.

We tested how sizes of deletions and genomic locations affect Swap-seq editing efficiency. We selected a candidate enhancer regulating *PPIF* and chose pegRNAs that have shown high editing efficiency from a previous work (**table S1**). We designed pegRNA pairs to delete 55 to 215 bp, transfected HEK293T cells, amplified the genomic sequence, and visualized editing outcome through gel electrophoresis.

### Testing Swap-seq twinPE at endogenous loci in THP-1 cells

We designed Swap-seq barcode cassettes (**table S2**) based on experimental evidence (**fig. S1**, see **Swap-seq pegRNA optimization in HEK293T cells**) and computational predictions (**fig. S2**, see **Barcode design guided by ChromBPNet predictions**). To test Swap-seq editing efficiency and accuracy in regions with different chromatin accessibility and histone modifications, we designed pegRNA pairs that delete from 71 to 215 bp within three regions: the promoter of *PPIF*, a candidate enhancer, and a region with low ATAC-seq signal (**table S1**, **Fig. 1B**). We infected dox-inducible PE2-THP1 cells with lentivirus carrying pegRNA pairs cloned on a single plasmid (see **Cloning Swap-seq pegRNA pairs and pegRNA library**) at low MOI (0.3), added doxycycline for 2 weeks, and extracted genomic DNA for PCR amplification. We designed PCR primers with 15-bp unique molecular identifiers (UMIs) to avoid PCR/sequencing bias as edited alleles are significantly shorter than WT. After 3-round PCR as previously described ^40^, amplicons were sequenced on an Illumina sequencing platform. We deduplicated the sequencing read with AmpUMI ^114^and analyzed the data with CRISPResso2 ^113^.

### Swap-seq *PPIF* tiling deletion screen design

#### Selection of candidate regulatory elements at the PPIF locus

We selected candidate regulatory elements by integrating public epigenetics data in THP-1 and our previous CRISPRi-FlowFISH results at this locus^45^. We selected candidate regulatory sequences from 5 major categories: (1) Four candidate distal enhancers that significantly decreased *PPIF* expression when targeted with CRISPRi; (2) 6 additional candidate enhancers that are accessible, have high levels of H3K27ac, and predicted by ENCODE-rE2G model^31^ as enhancers regulating *PPIF*, 3 of which are within the gene body and 3’ UTR of *PPIF*; (3) 5 accessible peaks that overlap with CTCF ChIP-seq peaks; (4) 1 accessible peak that had a small positive effect on *PPIF* (+8%, p=0.004) when targeted by CRISPRi; and (5) 8 additional intergenic peaks with lower accessibility and minimal H3K27ac. As several candidate regulatory elements are within the introns of the two nearby genes *ZMIZ1* and *ZCCHC24*, we also designed edits against their promoters as controls for *trans*-effects.

#### Negative controls: Barcode-only constructs

To calculate the effects of each edit, we need to measure the distribution of WT cells after RNA FlowFISH and cell sorting (see **Analysis of Swap-seq screens)**. Since unedited cells do not have barcode cassettes that could be amplified and sequenced, we designed lentiviral constructs with 6 different barcodes that only have the Swap-seq barcode cassette but without pegRNAs (**table S2**). We combined the barcode-only constructs with Swap-seq pegRNA library at 15% of the molecular weight of the pool. We made lentivirus and infected THP-1 cells at MOI of 0.3. After FlowFISH, we amplified the barcodes and used the distribution of 1 of the 6 barcodes associated with negative control constructs as the WT distribution, and the other 5 negative control barcodes to control for noise in the experimental process independent of editing (fig. S3B).

#### Negative controls: Out-of-chromosome edits

To distinguish genuine cis-regulatory effects from artifacts of the Swap-seq editing and RNA FlowFISH process, we included negative control pegRNA pairs that delete accessible peaks on different chromosomes from *PPIF*, which we referred to as out-of-chromosome edits. They are analogous to “safe-targeting” controls in CRISPR knockout screens^115^. Ideal negative control regions should allow high editing efficiency and have no regulatory effects on local gene expression to avoid potential trans-effects on the gene of interest. We selected *AAVS1*, a known safe-harbor locus, and *HEK3*, a locus with known high-efficiency gRNAs^40^. To identify additional accessible peaks that should have minimal effects on expression of local genes, we analyzed THP-1 CTCF ChIP-seq data^48^ to identify peaks that are at least 500 kb away from the closest protein coding genes, which we used as targets for out-of-chromosome control edits.

### Swap-seq pegRNA pairs design for high-throughput screens

#### Spacer design for Swap-seq deletion pairs

For each candidate regulatory element, we extended the coordinates of the target peak by 200 bp on each side to allow for additional gRNA choices, then used CRISPick ^116,117^ in CRISPRko mode (Quota = 500) to generate candidate SpCas9 (NGG, Chen(2013) tracrRNA) spacer sequences across the region. We removed low-quality candidates using a set of filters: (1) spacers containing an ambiguous (N) base, (2) spacers containing homopolymer stretches of more than four ‘‘T’’ nucleotides in a row (encoded as ‘‘U’’ in the RNA) that could prematurely terminate Polymerase III transcription of the pegRNA, (3) spacers with unbalanced GC content outside 20–90%, and (4) spacers with low-complexity sequences, such as 5-mer mononucleotide repeats. We then scored the remaining spacers for on-target efficiency (Rule Set 3) and specificity (cutting frequency determination (CFD) score) using FlashFry^118^, discarding spacers with a CFD specificity score below 0.2, a cutoff previously shown to minimize off-target effects^17^, or an on-target score below −0.5, chosen to retain sufficient guide density for pairing while still enriching for efficient spacers. We then removed any pegRNA pairs containing internal BsmBI or PaqCI recognition sites, which would interfere with cloning the library.

To generate candidate twinPE deletion pairs, we identified all remaining spacers with a cut-to-cut distance of 50–250 bp. Each region normally has more candidate pairs than we can select, because we want to optimize the library size to facilitate RNA FlowFISH and FACS sorting. Empirically, at coverage of 10,000 sorted cells per edit, we had >80% power to detect an effect size of 10% ^41^. Assuming 5% average editing rate and 20% cells surviving selection at MOI of 0.3, a library of 200 edits requires infecting at least 200M cells per biological replicates. We further prioritized deletion pairs to maximize the coverage of the whole region then included overlapping pairs so each deleted region has more than one deletion tile for cross-validation (see **Cloning Swap-seq pegRNA pairs and pegRNA library**). For each selected spacer, we designed the PBS to a fixed length of 11 nt, as previously described in ^41^.

#### Barcode cassette design guided by ChromBPNet predictions

To avoid creating TFBS within the barcode cassette by chance, we screened candidate cassettes *in silico* with the THP-1 ChromBPNet models^47^. We fixed the 21-bp overlap sequence and used the model to evaluate three candidate PCR primer binding sites that do not appear elsewhere in the genome: (1) adapted from Illumina adapters: CACGACGCTCTTCCGATCT /TCGTATGCCGTCTTCTGCTTG; (2) adapted from published MPRA design^119^: GTCACAAGGGCCGGCCACAA / AACTCCAGCAGGACCATGTGATC; (3) adapted from STARR-seq^26^: AGGCTTAAGCATGGCTAGCAAAG / GCCGCTTCGAGCAGACATG. For each architecture we enumerated the complete barcode space — all 4^7^ = 16,384 possible 7-mers, each followed by a fixed C. We inserted the barcode cassettes into twenty different background sequences generated by dinucleotide shuffles of an endogenous DNA sequence to generate 2,114-bp input sequences. We scored each sequence with the counts head of the bias-corrected ChromBPNet model^47^ trained on THP-1 ATAC-seq. Predictions were averaged across five-fold validation folds to yield a single predicted activity score.

The median predicted activities of each primer pairs are 3.966 (3.517–5.388) for the Illumina primers, 3.987 (3.418–5.808) for the MPRA primers, and 3.666 (3.433–5.123) for the STARR-seq primers. We chose STARR-seq primers because of their clean PCR amplification results and median predicted activity closest to the background sequence of 3.5. We then generated base-resolution sequence contribution scores using the DeepSHAP/DeepLIFT algorithm^120^ (see **ChromBPNet predictions, DeepSHAP sequence attributions and TF-MoDISco analysis**) and noticed that sequences with unusually high or low predicted activities were predicted to create TFBS by chance (fig. S2B). Therefore, we first chose 5,000 barcodes with predicted activities closest to the median activity, discarded any sequence that can be digested by BsmBI or PaqCI restriction enzymes, and greedily accepted barcodes if they have Hamming distance ≥2 from accepted sequences ranked by deviation from the median score. We divided the remaining 1,197 barcodes into 4 tiers based on their deviation from the median and assigned a random barcode from each tier to every Swap-seq edit.

Experimental results confirmed that this approach generated barcodes that have close to neutral activities, as Swap-seq effects from 4 barcodes are internally consistent, showing no particular barcode-associated artifacts (fig. S2G).

### Cloning Swap-seq pegRNA pairs and pegRNA library

We developed a three-round cloning approach based on previous work cloning dual Cas9 gRNAs to encode two paired pegRNAs on a single plasmid^36^.

In round 1, we assembled Multiplex Gene Fragments (ordered from Twist Bioscience, **table S2**) into a BsmBI-digested (New England Biolabs #R0739S), gel-purified (New England Biolabs #T1120S) lentiviral backbone (sgOpti_tevopreQ1, Addgene #255532). Gibson assembly was performed for 1 hour at 50°C using 20 uL of NEBuilder HiFi assembly (New England Biolabs #E2621L), 500 ng digested backbone, and 126.2 ng gene fragment pool in a 40 µL reaction, along with a negative control that does not have the gene fragment insert. We purified the Gibson assembly with 0.7X Ampure XP SPRI and eluted in 11 uL, then electroporated 10 uL into Endura competent cells (Lucigen #602422) using the 1 mm cuvette with the Bio-Rad micro pulser electroporation system (1.8 kV, 600Ω, 10 µF). We performed a serial dilution of 10 uL of the electroporated bacteria and spread it on agar plates containing carbenicillin, and expanded the rest in liquid culture containing carbenicillin for 20 hours at 30°C, then the library plasmid was extracted and purified with the ZymoPURE II Plasmid Midiprep Kit (Zymo Research #D4201). Success of Gibson assembly was assessed by counting colonies on the agar plate to ensure >1000-fold coverage of the library, and >100-fold increase over negative control.

After round 1 cloning, the plasmids now contain complete pegRNA1, BsmBI cloning site (cgcgagagacgaggctcgtctcatgttg), spacer of pegRNA2, PaqCI cloning site (gtttcaccgcaggtgtcacgcacctgcctagctgc), and RT template of pegRNA2. We designed a gene fragment that contains truncated evoPQ1 and polyT ends of pegRNA1, a 47-bp buffer sequence, and mU6 promoter for pegRNA2, flanked by BsmBI cloning sites at the ends. We seamlessly integrated this fragment into round 1 plasmids through a Golden Gate Assembly reaction, using 75 ng of Round 1 plasmids, digested with BsmBI and gel purified (we found digesting the plasmid prior to Golden Gate assembly improved efficiency and decreased background), 23.04 ng of the gene fragment, 2 ul of T4 DNA Ligase Buffer (10X), and 1 uL of NEBridge Golden Gate Enzyme Mix (BsmBI-v2) in a 20 uL reaction. Normally, 8 replicates of such assemblies were made in parallel. Each reaction uses the following incubation settings: 42°C 5 minutes, 16°C 5 minutes for 65 cycles, followed by 55 °C for 30 minutes and 80 °C for 20 minutes. We pooled the 8 replicates together and purified them with 0.7X Ampure XP SPRI, eluted in 11 uL, and electroporated 10 uL into Endura competent cells.

After round 2 cloning, the plasmid lacks only the scaffold sequence for pegRNA2. We used a previously published mutated gRNA scaffold to mitigate potential lentiviral recombination^121^. We chose not to encode the scaffold sequence for the second pegRNA because it will make the gene fragment too long and repetitive, increasing the chance for *E. coli* recombination and production failure. Similar to round 2, we integrated the scaffold using Golden Gate assemblies with 75 ng of pre-digested and gel-purified round 2 plasmid, 7.38 ng of the gene fragment, 5 uL of NEBridge Ligase Master Mix (NEB #M1100), 1.25 uL of PaqCI (NEB #R0745), and 0.625 µl of PaqCI activator (20 µM) in a 15 ul reaction. We prepared 8 replicates of the assembly using the following incubation settings: 37°C 5 minutes, 16°C 5 minutes for 65 cycles, followed by 37°C for 30 minutes and 65°C for 20 minutes. We pooled the 8 replicates together and purified them with 0.7X Ampure XP SPRI, eluted in 11 uL, and electroporated 10 uL into Endura competent cells. To sequence-validate pegRNAs and ascertain their relative abundance for use in Swap-seq analysis, we PCR amplified the pegRNAs from the final plasmid library and sequenced amplicons with Illumina MiSeq 500-cycle kit.

We followed the similar procedures for cloning individual plasmids that encode the pegRNA pair. Different from the pooled cloning protocol, in round 1, instead of transforming 10 ul of purified assemblies into Endura cells, we instead transformed 1 uL of the Gibson assembly product into 50 uL of NEB Stable Competent E. coli (New England Biolabs #C3040H), cultured overnight on agar plates containing antibiotics, and picked colonies for miniprep the next day. For rounds 2 and 3, one assembly reaction was enough and similarly, 1 uL of assembly product was used for transformation followed by plating and miniprep.

### Lentivirus production

We plated 550,000 HEK293T cells on 6-well plates coated with 1:20 dilution of Poly-D-lysine solution (Millipore Sigma #A-003-E) and 24 hours later we transfected the cells with 900 ng psPAX2 packaging vector (Addgene #12260), 360 ng pMD2.g VSV-G envelope vector (Addgene #12259), and 1.2 ug of purified plasmid library using 5.8 uL of X-tremeGENE HP^TM^ DNA Transfection Reagent (Millipore Sigma #06366236001). We harvested viral supernatant 48 hours later with 0.45 µm filtration. Harvested virus was used immediately or stored at -80°C.

### Lentiviral infection, selection, and doxycycline-induced prime editing

We transduced THP-1 cells harboring doxycycline-inducible PE2 at a multiplicity of infection (MOI) of 0.3 at a coverage of >80,000 cells per pegRNA or >320,000 cells per deletion tile. We chose low MOI to avoid multiple edits in a given cell. We transduced cells at high coverage to account for variable editing efficiency as we observed editing rate can be different by >100-fold between pegRNAs^41^. We titered each pegRNA library through a small scale transduction by adding serial volumes of lentivirus (0, 1, 3, 5, 10, 20, 50 uL) to 500,000 cells resuspended in 1 mL of complete RPMI media and 8 ug/mL of polybrene (Millipore Sigma #TR-1003-G) in 48-well plates, which we then centrifuged at 1,200 x g for 40 minutes at 37°C and cultured the infected cells in the incubator for 48 hours. We then performed 72 hours of selection using 1.5 ug/mL of puromycin, after which we measured viability of cells using Trypan blue and selected the volume of virus corresponding to 30% of cells alive. We transduced the pegRNA library using the same procedure and scaled the number of cells and amount of virus to reach the desired coverage. After the 72 hours of puromycin selection, we pelleted cells and resuspended in complete RPMI media containing 1.0 ug/mL doxycycline to induce prime editing and 0.3 ug/mL puromycin for maintenance selection. We maintained cells in media containing doxycycline for 14 days, then removed doxycycline and continued to maintain cells for 7 days before proceeding to downstream assays to ensure that PE2 was degraded and absent from the cells.

### PrimeFlow RNA FlowFISH assay

To measure the effects of Swap-seq edits on *PPIF* gene expression, we used the PrimeFlow RNA Assay Kit (Thermo Fisher Scientific #88-18005-210), similar to our previous work conducting *PPIF* FlowFISH in combination with CRISPRi or pooled prime editing^41,45^. Briefly, we used 12-15 million cells per reaction (5-7 reactions, referred to as FlowFISH replicates, and 1 control reaction per biological replicate) and performed the fixation and permeabilization steps in bulk for each biological replicate to reduce cell loss. The control reaction is used to assess background fluorescence signals from non-specific binding of the label probes (see **Cell sorting**). Target probes are as follows: gene of interest (*PPIF*; Thermo Fisher Scientific ‘Type 1’ probe set #VA1-30000735) and positive control housekeeping gene (*RPL13A*; Thermo Fisher Scientific ‘Type 4’ probe set #VA4-13187). Each sample (including the control reaction) was stained with the fluorescent Label Probes (Alexa Fluor 647 and 488 for *PPIF* and *RPL13A*, respectively) and we added a total of five washes with 40°C wash buffer following the final wash described in the manufacturer’s protocol to remove as much unbound label probe as possible before proceeding to cell sorting.

### Cell sorting

We sorted the cells following similar procedures as described in Variant-EFFECTS^41^. Briefly, we sorted 12-15M cells per FlowFISH replicate into 4 bins based on the fluorescence intensity of the target gene using the MA900 Multi-Application Cell Sorter (Sony). To control for differences in staining efficiency of each cell, we normalized the fluorescence associated with the gene of interest (*PPIF*) to that of a housekeeping control gene (*RPL13A*). Specifically, we used color compensation to subtract a portion of each cell’s AF647 signal (*PPIF*) based on the intensity of its AF488 (*RPL13A*) signal. This portion is selected such that the mean of the AF488 signal in the top and bottom 25% of cells based on AF647 was within 10% of the mean of the total sorted population’s AF488 signal. We set gates for each bin on the compensation signal to capture 25% of the cells across the full distribution. After sorting, we promptly stored the cells at 4°C.

### Genomic DNA extraction and barcode amplicon sequencing

We followed the exact protocol for genomic DNA extraction as described in our previous work^41^. Following extraction, we split genomic DNA into PCR replicates and amplified the Swap-seq barcode cassettes with Q5 High-Fidelity 2X Master Mix (New England Biolabs #M0492L) and 0.5 uM of each amplification primer (**table S1**) with the following program: 98°C for 1 minute and then 26 cycles of 98°C for 10 seconds, 69°C for 20 seconds, 72°C for 20 seconds, followed by a final 72°C extension for 2 minutes. We purified PCR products using a 1.5X SPRI bead clean-up and used 3 uL of the clean product as input for a secondary sample indexing PCR. Specifically, each 10 uL reaction contained 3 uL of sample, 5 uL of Q5 High-Fidelity 2X Master Mix, and 0.5 uM of each indexing primer containing Illumina adapters: CAAGCAGAAGACGGCATACGAGATNNNNNNNNGTGACTGGAGTTCAGACGTGTGCTCTTCCGATC T; AATGATACGGCGACCACCGAGATCTACACNNNNNNNNACACTCTTTCCCTACACGACGCTCTTCCG ATCT.

We carried out indexing PCR with the following thermocycling parameters: 98°C for 1 minute and then 10 cycles of 98°C for 10 seconds, 68°C for 20 seconds, 72°C for 20 seconds, followed by a final 72°C extension for 2 minutes. We pooled equal volumes of indexing PCR product together and performed 1.2X SPRI bead clean-up followed by another 1.0X SPRI bead clean-up, then measured DNA concentration using the Qubit 1X dsDNA HS Assay Kit (Invitrogen #Q33231). We sequenced DNA amplicon libraries on an Illumina sequencing platform.

### Analysis of Swap-seq screens

#### Sequencing, alignment and barcode counting

We demultiplexed raw reads with bcl2fastq (--use-bases-mask Y*,I8N2,I8N2,Y* --barcode-mismatches 0), trimmed sequences to the full 71 nt barcode cassette, and aligned them to barcode cassette references with CRISPResso2^113^ (CRISPResso -r1 {read1} -r2 {read2} -o results/crispresso/ --amplicon_seq {cassette} --amplicon_name Swap-seq_Barcode --name {SampleID} --exclude_bp_from_left 0 --exclude_bp_from_right 0 --plot_window_size 0 --min_frequency_alleles_around_cut_to_plot 0.05 --needleman_wunsch_gap_open -200 -q 20 -s 0). Because each edit-specific barcode cassette differs from each other only at the barcodes, CRISPResso2’s default gap-open penalty allowed the aligner to open gaps within the barcode cassette, resulting in misalignment. We therefore set the Needleman–Wunsch gap-open penalty to −200, which forces ungapped alignment across the barcode cassette. We designed barcodes with Hamming distance ≥2 with each other, so no single substitution error converts one designed barcode into another. Despite the rational design to avoid TFBS within barcode cassettes (see **Barcode cassette design guided by ChromBPNet predictions**), mutations could in theory create TFBS by chance. Therefore, we further filtered the aligned reads to assign each read to a barcode cassette by exact string matching. We allowed mutations outside of the barcodes for the non-editing barcode-only negative controls used as WT alleles, because mutations on these inert constructs will not affect gene expression.

#### Maximum-likelihood effect estimation

We summed the reads across PCR replicates within each FACS bin, and normalized these aggregated counts so that the sum of reference and variant alleles in each sample equals the total number of cells sorted into that FACS bin (i.e., to account for differences between the number of sequencing reads and the number of cells in a given bin). We used the limited-memory Broyden–Fletcher–Goldfarb–Shanno algorithm maximum-likelihood estimation (MLE) method in the R stats4 package to fit the read counts in each fluorescence bin to the log-normal distribution that has the highest likelihood for having produced the observed counts in the bins. To calculate effect sizes, we determined the percent difference between the mean of the log-normal fit for each edit and the mean of the log-normal fit for the reference barcode.

#### Heterozygous correction for Swap-seq

We then applied heterozygous correction to the results of MLE to estimate the true effects of the edits. Assuming independent transcription from the two alleles, the apparent expression from a heterozygously edited cell will be half the effect of a homozygously edited cell, leading to underestimation of the true effect sizes. To correct the estimated effect size, we need to measure the allele frequency to estimate the heterozygous rate for each edit. However, Swap-seq only reads out cells that are edited or carry barcode-only cassettes, thus the raw allele frequency of each edit we observed from the screen was that among cassette-bearing alleles instead of among all alleles. To solve this problem, we designed the barcode-only cassettes to serve dual purposes: (1) we used the distribution of barcode-only cassettes among expression bins as the reference distribution (see Negative controls: Barcode-only constructs). (2) We added the barcode-only cassettes to the plasmid library at 15% of the molar ratio. We assumed all cells that received the barcode-only cassette have hemizygous integration of the barcode cassettes, thus after antibiotic selection, barcode-only cassettes should constitute 7.5% of all alleles. We then calculated the allele frequencies *f_i_* of each edit using the following equation:

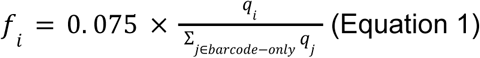

where *q_i_* is the observed frequency of allele *i* among cassette-carrying alleles. We genotyped edited loci and confirmed the accuracy of this method (**fig. S4E**). We applied heterozygous correction as previously described in Variant-EFFECTS with modifications for Swap-seq.

#### Terms

*f_i_* = allele frequency in all cells

*x_i_* = pegRNA (‘‘guide’’) frequency in plasmid pool, assuming this is the same in the infected cells

*g_i_* = edit frequency in cells that carry guide *i* (‘‘editing efficiency’’)

*e_i_* = true effect size of homozygously edited cell (*i*/*i*), this is defined such that *e*_0_ = 0 and a value of e.g. -0.2 means that *i*/*i* cells have 80% as much mRNA as a homozygous wild-type cell

*m_i_* = measured effect size of edit (in FlowFISH space)

*m_o_* = WT gene expression

*n_i_* = normalized effect size, after dividing by wild-type effect (*m_i_*/*m_o_*)

#### Genotype frequencies assuming independent editing on each allele

0/*i* (ref/alt) = 2*x*_i_*g_i_*(1 − *g*_i_)

*i*/*i* (alt/alt) = *x_i_g_i_*^2^

We assume that the true effect size of a diploid cell with alleles *i* and *j* (*e_ij_*) is given by averaging the effects of the two alleles, converted to fluorescence units from FACS (*Z*):

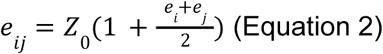

and that the observed effect of a single variant (*m_i_*) is a weighted sum of the effect sizes and frequencies of each genotype *j* containing that allele.

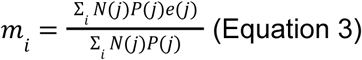

Here *N*(*j*) represents the number of times allele *i*appears in genotype *j*, *P*(*j*) represents the frequency of the genotype, and *e*(*j*) represents the effects of genotype *j*. For non-WT edits, *m_i_* can be calculated

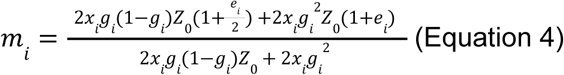

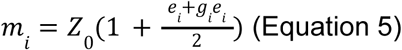

As the MLE-estimated effects were computed in FlowFISH space, we normalize the effect of each allele *i* by that of the wild-type:

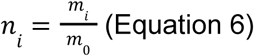

Swap-seq uses the distribution of non-editing barcode-only cassettes as the reference distribution, which we assume have the same distribution as homozygous WT cells. Therefore, the observed WT gene expression is the fluorescence background *Z_0_*, plugin into Equation 6:

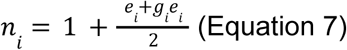

Then, the true effect size (*e_i_*) of homozygously edited cell is simply:

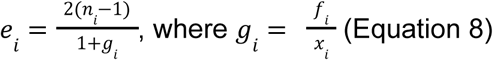

We measured *x_i_* through sequencing the plasmid library, estimated *f_i_* FlowFISH library (Equation 1), and calculated *n_i_* using the results of MLE. through sequencing the RNA

#### Setting frequency filters by assessing reproducibility

Allele-frequency thresholds were evaluated by their effect on replicate reproducibility and on the number of edits retained (**fig. S4B**). Without a threshold, correlation between replicates is poor (median Pearson’s r = 0.35 for pairwise FlowFISH replicates; Pearson’s r = 0.69 for biological replicates). Edits at low allele frequency are supported by few reads, so their effect estimates are dominated by sampling noise. Reproducibility rose with the threshold and saturated at the allele frequency of 10^-5^, beyond which further increases discarded many edits with minimal improvement of correlation (<0.05 increase of Pearson’s r).

A barcode was therefore required to reach an allele frequency of 10^-5^ in any given FlowFISH replicate. To account for barcode-to-barcode variation, we required each edit to have ≥ 3 barcodes pass in at least one FlowFISH replicate of each biological replicate, unless 2-barcode support already yielded a significant effect (see *Quantification and statistical analysis*). Two of the FlowFISH technical replicates in biological replicate 1 (replicates 4 and 5) were excluded from downstream analysis due to experimental errors. Effect sizes were calculated from the remaining technical replicates across both biological replicates. Edit effects were calculated by averaging barcode effects within each FlowFISH replicate, then averaging those values across replicates.

#### Quantification and statistical analysis

The significance of each Swap-seq edit effect was assessed by a two-sided one-sample t-test of its per-replicate effects against a null of no change, and p values were corrected using the Benjamini-Hochberg procedure. Edits with a BH-corrected p < 0.001 were called significant. A regulatory element was called significant if at least one of the edits tiling it was significant.

### Single-edit induction and clonal cell line selection

We infected dox-inducible PE2 THP-1 cells with a pair of pegRNAs following the same procedure as described before (see **Lentiviral infection, selection, and doxycycline-induced prime editing**). After 2 weeks of doxycycline treatment to induce prime editing, we sorted cells into U-bottom 96-well plates to generate single clone cell lines through FACS. After 1 month of expansion, we extracted gDNA from 250,000 cells from each clone using QuickExtract DNA Extraction Solution (BioSearch Technologies #QE09050), following manufacturer’s instructions. We then amplified the target locus using PCR and sequenced on an Illumina platform. Homozygous wild-type and homozygously edited cells were defined as having <10% or >90% of reads mapping to the desired genotype, respectively^41^. For experiments associated with **fig. S4C**, we used a simplified protocol to select homozygous WT and homozygous edited clones through gel electrophoresis as the deletion sizes are larger than 100 bp and PCR products of WT and edited alleles can be clearly separated on a gel.

### RNA extraction and quantitative RT-PCR

We expanded clonal lines in 24-well plates and extracted total RNA from 1M of each clone with Quick-RNA Miniprep Kit (Zymo Research R1055) according to the manufacturer’s instructions. We measured the concentration and purity of all extracted RNA samples with a NanoDrop. For each RT-qPCR reaction, we added approximately 200 ng of total RNA and 400 nM of each primer (**table S1**) to 10µL of Luna Universal One-Step Reaction Mix and 1 µL of Luna WarmStart RT Enzyme Mix (New England Biolabs #E3005E) in a 20 µL reaction. We used the following program: 55°C for 10 minutes for cDNA generation, 95°C for 1 minute, followed by 45 cycles of 95°C for 10 seconds, 60°C for 30 seconds with fluorescence detection, followed by a melt curve step. We calculated the effects of each clonal line using the ΔΔCT method by comparing the housekeeping-normalized CT of WT and edited clones.

### Cloning downstream lentiMPRA

To test the gene regulatory function of the candidate CREs and avoid potential confounding effects, such as transcription initiation from the CREs, we previously developed the downstream MPRA, where the candidate CREs were placed after the termination sequence of the reporter gene^50^. Because THP-1 cells are hard to transfect^122^, we cloned the downstream MPRA sequence into a lentiviral construct^68^. We cloned the plasmids individually to avoid lentiviral recombination causing mismatches between promoters, candidate CREs, and barcodes in the reporter transcript which we sequenced to calculate the CRE activity scores^123,124^. Since the reporter construct contains a splice site, we cloned the MPRA sequence in the reverse orientation relative to the CMV promoter that transcribes the lentiviral RNA to avoid aberrant splicing of the reporter transcripts.

To clone the downstream lentiMPRA construct, we first digested the lentiviral backbone (**table S4**) with AgeI (NEB #R3552) and EcoRI (NEB #R3101), followed by gel purification. We then prepared four other fragments for a Gibson assembly reaction: (1) we amplified the promoter fragments to add Gibson assembly handles, using 30 µL of Q5 High-Fidelity 2X Master Mix (New England Biolabs #M0492L), 0.5 uM of each primer, and 1 ng of the gene fragment in a 60 µL reaction (98°C for 30 seconds, and 11 cycles of 98°C for 15 seconds, 66°C for 15 seconds, 72°C for 60 seconds, followed by a final 72°C extension for 2 minutes). We then purified the PCR products with a 1.2X SPRi clean. (2) We designed five 8 bp barcodes for each promoter-CRE pair and added these barcodes to the ORF fragment through a 2-round PCR process. In round 1, we combined 25 µL of NEBNext Ultra II Q5 Master Mix (New England Biolabs #M0544L), 1 ng of the hSTARR-seq_SCP1 (Addgene Cat# 99319), 10 µM of the forward primer (GGCCCGGTCGTGAG), and 10 µM of a pool of five reverse primers, each carrying a specific barcode (GCCGCCCCGACTCTAGNNNNNNNNTGCATGCTCTAGATCAATCTAATTCAAC), in a 50 µL reaction (98°C for 30 seconds, and 7 cycles of 98°C for 10 seconds, 65°C for 20 seconds, 72°C for 30 seconds, followed by a final 72°C extension for 2 minutes). After a 1X SPRi clean, we eluted the PCR1 product in 20 µL of water and used 1 µL for another PCR to add Gibson assembly handles by adding it to 25 µL of NEBNext Ultra II Q5 Master Mix and 0.5 µM of each primer (Fwd: GGCCCGGTCGTGAG, Rev: gtatcttatcatgtctgctcGAAGCGGCCGGCCGCCCCGACTCTAG) in a 50 uL reaction (98°C for 30 seconds, and 10 cycles of 98°C for 10 seconds, 67°C for 20 seconds, 72°C for 30 seconds, followed by a final 72°C extension for 2 minutes), followed by a 1X SPRi clean. (3) We designed the SV40 polyA sequence and a buffer sequence between the reporter transcript and candidate CRE as a gene fragment that can be directly used (“ds pAS spacer v2.2”). (4) For candidate CREs, we used the coordinates of the corresponding ATAC-seq peaks to fetch the sequences from hg19 reference genome, designed gene fragments, and PCR amplified to add Gibson Assembly handles, using 30 µL of Q5 High-Fidelity 2X Master Mix (New England Biolabs #M0492L), 0.5 uM of each primer, and 1 ng of the gene fragment in a 60 µL reaction (98°C for 30 seconds, and 11 cycles of 98°C for 15 seconds, 59°C for 15 seconds, 72°C for 60 seconds, followed by a final 72°C extension for 2 minutes). We purified the PCR product with a 1X SPRi clean.

For Gibson assembly, we added 0.05 pmol of the linearized plasmid and each prepared fragment to 10 µL of NEBuilder HiFi DNA Assembly Master Mix (New England Biolabs #E2621L) in a 20 µL reaction, followed by a 1-hour incubation at 50°C. We then transformed 15 µL of NEB Stable Competent E. coli (New England Biolabs #C3040H) with 2 µL of the Gibson Assembly product, cultured at 30°C for 20 hours, and extracted plasmids with miniprep. We confirmed cloning results and barcode association with whole plasmid sequencing.

### Downstream lentiMPRA experiment and analysis

We plated 350,000 HEK293T cells on 12-well plates coated with 1:20 dilution of Poly-D-lysine solution (Millipore Sigma #A-003-E) and 24 hours later we transfected the cells with 450 ng psPAX2 packaging vector (Addgene #12260), 180 ng pMD2.g VSV-G envelope vector (Addgene #12259), and 600 ng of individual plasmids using 2.8 uL of X-tremeGENE HP^TM^ DNA Transfection Reagent (Millipore Sigma #06366236001). We added 2µl ViralBoost (Alstem, #VB100) to each well of HEK293T cells 24 hours after transfection, harvested and pooled all viral supernatant the day with 0.45 µm filtration, and added LentiX Concentrator (Takara, #631232) to viral supernatant. After an overnight incubation at 4°C, we centrifuged the viral supernatant at 1,500 x g for 45 minutes at 4°C, removed supernatant, and resuspended pellets in complete DMEM, which was used immediately or stored at -80°C. We titrated lentivirus following published protocols^68^: we infected THP-1 cells with 0, 1, 2, 4, 8, 16, 32, or 64 µL lentivirus, and used qPCR to measure the copy number of viral integration. In our experiments, 32 µL of the lentivirus per 500,000 cells achieved high MOI without significantly affecting cell viability.

We infected 10M THP-1 cells for 2 biological replicates. 72 hours after spinfection, we extracted genomic DNA from 2M cells (Monarch Spin gDNA Extraction Kit, New England Biolabs #T3010L) and total RNA from 8M cells (Zymo Research #R1055) with DNase treatment following manufacturer’s protocol. We isolated polyA mRNA (NEBNext High Input Poly(A) mRNA Isolation Module, New England Biolabs #E3370S) and eluted in 18 µL of water. We then split each polyA mRNA into three technical replicates and treated each polyA mRNA sample with ezDNase (Invitrogen, #18091150) following the manufacturer’s protocol. We reverse transcribed RNA using SuperScript IV Reverse Transcriptase (Invitrogen, #18091150) and the gene specific primer DB31_STARR-seq_RT following the manufacturer’s protocol, using 9 µL of polyA mRNA treated with ezDNase and 1 µL for no RT controls (without using SuperScript IV Reverse Transcriptase) as a stringent control for genomic DNA contamination. After a 1.5X SPRi clean, we amplified cDNA with primers spanning the splice junction (**table S4**, DB032_Starr-jPCR_Fwd and DB033_Starr-jPCR_Rev) to further reduce genomic DNA contamination. We added all cDNA to 25 µL of NEBNext Ultra II Q5 Master Mix and 0.5 µM of each primer in a 50 uL reaction (98°C for 45 seconds, and 17 cycles (or 21 cycles for no-RT control samples) of 98°C for 15 seconds, 65°C for 30 seconds, 72°C for 30 seconds, followed by a final 72°C extension for 1 minute), followed by a 1.2X SPRi clean. We did a final round of PCR with amplified cDNA product and genomic DNA to add sample barcode and Illumina sequencing adapters using primers JR89/90 (**table S4**) at 0.5µM and annealing temperature of 69°C. The PCR product was then cleaned up using 1.2X SPRI beads and final product concentration was measured using Qubit HS dsDNA kit (Invitrogen #Q33231).

We sequenced the RNA and DNA library using Illumina NextSeq 500/550 with custom sequencing primer DB045_ExP_fwd_BC_seq for Read 1 (26 bp) and JR91_MPRA-RNA-Index1 for Index 1 (8 bp) (**table S4**). We demultiplexed the sequencing data and assigned each read to a barcode using matcha^125^, requiring ≤1 mismatch and unambiguous assignment. We averaged the barcode counts of DNA and RNA samples of 3 technical replicates and normalized by the total number of reads of each sample. For each barcode, we calculated the activity score as log2(normalized RNA / normalized DNA). We included a “noTFBS” synthetic sequence designed to be inactive in a previous study^50,126^ to measure the basal strength of each promoter. We then compared the activity scores of each candidate element to the basal strength of the respective promoter, i.e., log2 fold change = candidate element activity − basal, and percent change = (2^(candidate element activity − basal) − 1) × 100.

### Micro-C and RCMC data generation

Genome-wide Micro-C libraries were generated largely as previously described, with minor modifications^52,127^. For genome-wide THP-1 Micro-C, four independent biological replicates, totaling 80 million cells, were prepared from 20 million cells per replicate. For RCMC, three independently derived clones were analyzed for each condition, including WT and clonal lines carrying Swap-seq edits at CTCF c2 (see **table S2** for pegRNAs used). To reduce clonal variation, 40 million cells were used per condition in total.

Briefly, cells were sequentially crosslinked with disuccinimidyl glutarate (DSG) and formaldehyde (FA). Following an MNase titration experiment to determine the appropriate amount of MNase for digestion, chromatin was digested with MNase, DNA ends were repaired and labeled with biotinylated nucleotides, and proximity ligation was performed. Residual biotin was removed from unligated DNA ends, crosslinks were reversed, and DNA was purified. Dinucleosome-sized ligation products of 200–400 bp were isolated by agarose gel electrophoresis, enriched using streptavidin beads, and converted into indexed Micro-C sequencing libraries. For RCMC, custom biotinylated oligonucleotide capture probes from Twist Bioscience were designed to tile an approximately 2-Mb interval encompassing the *PPIF* locus. Indexed Micro-C libraries were subjected to hybridization capture, streptavidin pulldown, and post-capture amplification before paired-end sequencing.

### Micro-C/RCMC data processing

Micro-C and RCMC reads were processed using Snakemake^128^ workflows adapted from previously described Micro-C/RCMC pipelines^52,127^. Raw reads were quality-filtered with fastp^129^ using a qualified-base Phred threshold of 20 and a minimum read length of 30 bp. Reads were aligned to the hg19 reference genome with bwa-mem2^130^ in paired-end mode using the -SP option, and the alignments were streamed directly into pairtools^131^ parse2. Read pairs were converted to pairs format using the --add-columns mapq, --expand, --report-position outer, --drop-sam, and --assembly hg19 options. The --min-mapq 30 and --max-insert-size 150 options were applied during parsing.

For RCMC, parsed pairs were restricted to intra-region contacts for which both ends mapped within the capture-analysis interval, hg19:chr10:80,107,225–82,115,090. Genome-wide Micro-C data were processed analogously without this regional restriction. For each clone, pairs from all sequencing lanes and, where applicable, technical sequencing runs were coordinate-sorted and merged, followed by a single deduplication step with pairtools dedup using the --max-mismatch 1 and --mark-dups options. Deduplicated biological-replicate pairs were retained separately for replicate-level analyses and were additionally merged without further deduplication to generate pooled condition-level contact maps.

Deduplicated pairs were indexed with pairix^132^ and loaded into cooler^133^ at 50-bp base resolution using the hg19 chromosome sizes. Multi-resolution .mcool files were generated with cooler^133^ zoomify using the --balance option.

### RCMC data analysis

We performed analysis on the deep-sequenced RCMC dataset for wild-type (WT) and Swap-seq deletion clones at the *PPIF* locus. For each genotype, we used contact matrices derived from three clonal cell lines for all statistical comparisons and pooled matrices for map-level visualization, as indicated below.

We calculated insulation scores using the ‘cooltools’ package^134^ (version 0.7.1) on Python version 3.11.15. For each clone of WT or deletion genotype, we extracted contact matrices of chromosome 10 from .mcool files at 1 kb resolution into standalone coolers using the ‘cooler’ package (version 0.10.4), with pixels reindexed to chromosome-local bins. We then computed insulation scores with a window size of 50 kb. For each deleted site, the score was calculated from the 1 kb bin containing the midpoint of the deletion and compared between the three WT clones and the three clones carrying that deletion.

To generate differential contact maps (**Fig. 2F**), we aggregated the per clone contact matrices of WT or CTCF deletions at 1.6 kb resolution, and extracted the data within a 1 Mb window centered on the *PPIF* promoter using the ‘cooler’ package (version 0.10.4) on Python version 3.11.15. We calculated the change of contact per pixel between deletion and WT as log2((deletion + pseudocount) / (WT + pseudocount)), smoothed with a 3 x 3 mean filter.

To quantify differential contacts between regulatory elements, we extracted ICE-normalized contact matrices (MAPQ≥30) at 1, 1.6, 2 and 3.2 kb resolution from per-clone .mcool files using the cooler package (version 0.10.4) on Python version 3.11.15. For each clone, we quantified contacts between a viewpoint (i.e. *PPIF* promoter or silencer s1) and each target regulatory element as the mean of a 3 x 3 pixel block, defined by taking the bin containing each feature’s midpoint, expanding it by one bin in either direction, and intersecting the two resulting three-bin intervals. We added a pseudocount, defined as the 1st percentile of positive block pixels pooled across genotypes divided by ten, to each block mean. We calculated log2 fold change as the difference between the mean log2 block contact of the three deletion clones and that of the three WT clones, and assessed significance with an unpaired, two-sided Welch’s t-test.

### Fluorescent reporter assay and flow cytometry

We cloned the fluorescence reporter constructs by first moving a previously published mCitrine reporter^74^ into a lentiviral backbone, and then changed the promoter and upstream regulatory sequences. Briefly, to change the promoter sequence, we digested the plasmid with NsiI and BstXI, followed by Gibson Assembly using 0.05 pmol of linearized plasmid, 0.1 pmol of gene fragment, and 10 µL of NEBuilder HiFi assembly in a 20 µL reaction. Similarly, we digested the plasmid with SnaBI and NsiI followed by Gibson assembly to replace the upstream 9xTetO sequence with regulatory elements. We then infected WT THP-1 cells with lentivirus carrying fluorescent reporter in 3 biological replicates, treated cells with 1.5 ug/mL of puromycin to select for infected cells, and measured mCitrine fluorescence with flow cytometry analysis (Bio-Rad ZE5, Everest v.2.3-3.0). For chemical inhibition to analyze cofactor dependencies of silencer s1, we added 10 μM TMP269 (Selleck Chemicals #S7324) or 10 μM tazemetostat (Selleck Chemicals #S7128) to culture media on Day 0 to inhibit class IIa HDACs or EZH2, respectively. Drug conditions were matched with a DMSO vehicle control and mCitrine expression of constructs containing silencer s1 upstream of the promoter was compared to matched promoter-only controls for each drug.

### 5’-end TSS mapping

To sequence the 5’-end of reporter transcripts and capture potential transcriptional activities from the silencer s1, we obtained purified polyA+ mRNA, reverse transcribed it with a RT primer and a template switch oligo (TSO), and sequenced with a MiSeq 150-cycle kit as previously described^50^. We modified the RT primer sequence to be closer to the promoter and silencer to capture potentially unstable transcripts (**table S1**). We demultiplexed sequencing reads by dual 8-bp index reads (bcl2fastq, exact index match, no mismatches allowed), yielding two biological and two technical replicates per reporter construct. We trimmed R1 reads to remove the TSO sequence (AAGCAGTGGTATCAACGCAGAGTACAT) from the beginning and the Illumina adapter from the 3′ end, and aligned trimmed R1 reads with bowtie2^135^ in --very-sensitive-local mode to a custom reference sequence spanning 2 kb upstream of the reporter start codon through 100 bases past the RT primer binding site. For constructs with the *EF1A* promoter, we aligned the reads to an intron-deleted reference. We discarded unmapped reads, merged technical replicates, and quantified TSS frequency as the distribution of alignment start positions.

### ATAC-seq

We applied ATAC-seq on 4 WT clones and 6 clonal lines carrying homozygous deletion of the silencer s1 as previously described^45^. We then sequenced the libraries on an Illumina NextSeq 550 (2 x 40 bp, dual-indexed) and demultiplexed with bcl2fastq2. We processed the datasets using the ENCODE ATAC-seq pipeline v2.2.3 (https://github.com/ENCODE-DCC/atac-seq-pipeline) with the standard settings and automatic adapter detection, aligned reads to the GRCh37/hg19 reference genome using Bowtie2^135^, and identified peaks using MACS2^136^. For each clone, we supplied two technical replicates into a single pipeline run to produce a pooled and reproducible peak set. All ten clones passed ENCODE quality thresholds (TSS enrichment 25.5–31.2; fraction of reads in peaks 0.47–0.63; naive-overlap and IDR reproducibility "pass"). We generated read-depth normalized coverage tracks from the pipeline’s alignments using deepTools bamCoverage (--normalizeUsing CPM --binSize 10 --extendReads --centerReads --ignoreDuplicates).

### CUT&RUN for histone modifications

We performed CUT&RUN on WT or clonal cell lines carrying silencer s1 deletions with the CUTANA ChIC/CUT&RUN kit (EpiCypher #14-1048) according to manufacturer’s instructions from manual version 6. We tested a range of Digitonin concentrations and determined that 0.01% digitonin was the lowest concentration that leads to complete permeabilization of THP-1 cells, which we used for all following experiments. We followed the standard protocol for CUT&RUN experiments targeting H3K27me3 (Antibody: EpiCypher 13-0055) and H3K4me3 (Antibody: EpiCypher 13-0060), using Rabbit IgG (Antibody: EpiCypher 13-0042) as a negative control. We used 0.5 µg of antibody for all experiments. We prepared the sequencing library using the CUT&RUN Library Prep Kit (EpiCypher 14-1001/14-1002) with 5 ng of input DNA according to the manufacturer’s instructions. We quantified DNA with Qubit and fragment sizes with Bioanalyzer, then sequenced the library at a targeted depth of 10M paired-end reads per sample on an Illumina NextSeq 550 with 2x75 cycles.

We optimized CUT&RUN conditions targeting H3K27ac and tested three antibodies (SKU: 13-0059, Active Motif 39133, and ab4729) with the standard protocol (no fixation), or in combination with two fixation conditions, following the manufacturer’s instructions (https://support.epicypher.com/docs/cross-linking-for-cut-and-run). Briefly, we resuspended 500,000 THP-1 cells in cold PBS, added formaldehyde to a final concentration of 0.1% (light fixation) or 1% (medium fixation), briefly vortexed, and incubated at room temperature for 1 minute. We then added glycine to a final concentration of 125 mM to quench the reaction, centrifuged at 600 x g for 3 min, decanted the liquid, and proceeded with the standard protocol for CUT&RUN, library prep and sequencing as stated above. ab4729 with light fixation generated the highest quality results among all conditions, as measured by clear enrichment of H3K27ac signal at TSSs genome-wide, which we used for all subsequent experiments.

To process CUT&RUN data, we adapted the previously published pipeline in https://zenodo.org/records/13698181^137^. We aligned paired-end reads to the human reference genome hg19 (male.hg19.fa) with Bowtie2^135^ v2.4.2 using --local --very-sensitive-local --no-unal --no-mixed --no-discordant --phred33 -I 10 -X 700, and to the *E. coli* genome with identical settings to quantify the spike-in DNA. We sorted and indexed the alignments with samtools^138^ v1.11 and removed duplicates with Picard^139^ v2.26.3 MarkDuplicates (--REMOVE_DUPLICATES true). For each library we calculated a spike-in scale factor as the reciprocal of the percentage of unique reads aligned to the *E. coli* genome: 1 / (100 × unique E. coli fragments / unique hg19 fragments). We then generated normalized coverage tracks with deepTools bamCoverage (--extendReads --centerReads --binSize 10 --scaleFactor).

### Differential signal analysis for ATAC-seq and CUT&RUN

We used a fixed set of ATAC-seq peaks generated from published bulk THP-1 ATAC-seq data for differential signal analysis so that identical intervals could be used across the ATAC-seq and CUT&RUN datasets. We filtered the MACS2 peaks called on bulk THP-1 cells to q < 0.05, coordinate-deduplicated, and extended each peak ±500 bp as the quantification windows. We counted fragments in each quantification window with featureCounts v2.1.1 from the Subread package (-p --countReadPairs -B -C -O -Q 10), including only properly paired, non-chimeric fragments with mapping quality ≥ 10, aggregating counts from the technical replicates of each clone. We tested differential accessibility or CUT&RUN signal between WT and silencer-deleted clones using pyDESeq2^140^ v0.5.4 ∼condition mode with DESeq2 median-of-ratios size factors and Cook’s-distance outlier refitting. We reported the log2 fold changes as the maximum-likelihood estimates from the negative-binomial Wald test and Wald p-values generated by DESeq2.

### Variant-EFFECTS screen at the s1 silencer

We targeted the *PPIF* silencer s1 identified in our Swap-seq screen, a 432-bp element at chr10:81,163,999–81,164,430 (hg19). Within this element we defined a 200-bp editing window (chr10:81,164,105–81,164,304) for prime editing, chosen so that all edits fall inside a single 247-bp PCR amplicon that can be sequenced on a NextSeq 550 300-cycle kit. We designed the pegRNAs for 87 edits that fall into three categories: (1) consecutive, end-to-end 5-bp deletions tiling the whole 200-bp window, (2) 5-bp mutations that overlap with 6 TFBS nominated by chromBPNet predictions, each replacing the WT sequence with three 5-bp scramble sequences (AGCCA, CAGTC, CTCAG) drawn from the pre-defined, GC-balanced substitution set used in Variant-EFFECTS^41^, and (3) *de novo* motif insertions. We designed and cloned the pegRNA library following the Variant-EFFECTS protocol ^41^. We generated lentivirus with the plasmid library, infected THP-1 cells at MOI of 0.3, and selected infected cells with puromycin. After 14 days of doxycycline treatment to induce prime editing, cells were cultured in doxycycline-free media for 7 days. We then measured effects of each edit with RNA FlowFISH, extracted and sequenced the genomic DNA from each expression bin, and analyzed the data using the Variant-EFFECTS pipeline (https://github.com/EngreitzLab/Variant-EFFECTS).

### ChromBPNet predictions and motif analysis

We used the five-fold ChromBPNet models trained on THP-1 ATAC-seq as described previously^41^. All values reported here are the average across the five-fold cross-validation of the bias-factorized model, taken from the counts head. To generate ChromBPNet predictions for each regulatory element, we defined a 2,114-bp input window per element, each centered on the span of deletion tiles targeting that element and extending to either side. We then passed the one-hot encoded DNA sequence of each window to the model, and calculated the fold-average model output as the predicted accessibility of that element.

To analyze base-resolution contribution to accessibility, we used the DeepSHAP/DeepLIFT^120^ algorithm as previously described^41^. Briefly, we created 20 dinucleotide-shuffled versions of the input sequences as reference sequences and then utilized the DeepSHAP implementation of DeepLIFT (https://github.com/kundajelab/shap) to generate sequence contribution scores. This yielded one 2,114 × 4 contribution matrix per element after averaging across folds. For display, contributions of each element were multiplied by the one-hot encoded DNA sequence so that each position carries only the contribution of the base actually present and its height scaled by the contribution score.

We annotated model-predicted transcription factor binding sites in each element using TF-MoDISco ^49^ and Fi-NeMo (https://github.com/kundajelab/Fi-NeMo). TF-MoDISco identifies recurring high-contribution patterns across a set of contribution-score profiles, clustering them into consolidated motifs summarized by a contribution weight matrix (CWM). We used the previously generated TF-MoDISco pattern set from the counts head of the THP-1 ChromBPNet model, which contains 57 patterns (51 positive, 6 negative), each a 30-bp contribution weight matrix (CWM) annotated by using TOMTOM from the MEME Suite^78^. We then used Fi-NeMo to call individual motif instances within each element. We passed the 2,114-bp DNA sequence of each element and its contribution matrix to Fi-NeMo using default parameters (-l/--global-lambda 0.7, -t/--cwm-trim-threshold 0.3), which returned a list of motif hits for each element, each assigned a transcription-factor identity from the top-ranked TOMTOM match of its TF-MoDISco pattern. Fi-NeMo’s default post-filter discards any hit whose cosine similarity to its CWM falls below λ, so every reported instance has similarity ≥ 0.7. Additionally, we assigned instances with hit importance ≥ 0.1 as high-confidence calls.

We preferred Fi-NeMo over a sequence-based scan such as FIMO^79^ because Fi-NeMo identified transcription factor motif hits using the attribution of the cell-type specific ChromBPNet model rather than the sequence alone. This approach takes into account the cellular (e.g. expression of TF and cofactors) and endogenous sequence (e.g. TF-TF interactions) context.

### Identification of other genomic silencers similar to s1

We analyzed candidate regulatory elements genome-wide to nominate candidate silencers based on the chromatin features of *PPIF* silencer s1. We defined candidate regulatory elements as ATAC-seq MACS2 narrow peaks, excluding elements annotated as promoters by a previous Activity-by-Contact (ABC) run on THP-1^45^. We further excluded accessible peaks that overlap with CTCF ChIP-seq peaks or peaks with ≥50% of the interval covered by repetitive sequences using RepeatMasker annotation. For remaining candidate elements, we used the ATAC-seq values in reads per million over the 500-bp candidate regions to quantify chromatin accessibility. To quantify H3K27ac signal, we extended the quantification windows to 2 kb centered on the midpoint of the ATAC-seq peaks and analyzed publicly available THP-1 H3K27ac ChIP-seq using pyBigWig.

We restricted the analysis to the most accessible elements by setting a threshold at the 80th percentile of ATAC-seq RPM (ATAC-seq ≥ 2.613 RPM). We then calculated a score for each peak as log2(H3K27ac signal over the 2 kb window / ATAC-seq RPM) and defined silencer-like and enhancer-like populations as peaks with bottom 15% (accessible but not acetylated) or top 15% (accessible, highly acetylated) of the log2 scores, respectively.

We assigned a nearest annotated gene to each silencer-like and enhancer-like peak as a proxy for its likely target, using the RefSeq collapsed gene-bounds annotation, same as the ABC pipeline^45^ (hg19 RefSeqCurated). We used previously published THP-1 RNA-seq data^45^ to assign a TPM value for each gene and compared the nearest gene expression of silencer-like and enhancer-like peaks with a two-sided Mann-Whitney U test.

We analyzed whether silencer s1 motifs identified by the Variant-EFFECTS screen and motif analysis, including SP1, GABPA, CEBPA, RUNX, ZNF324 and IKZF, were enriched in the silencer-like peaks vs. enhancer-like peaks. To avoid distance-related motif distribution confounding this analysis, we constructed sets of silencer-like and enhancer-like peaks with similar distances to the nearest gene, yielding 1,689 peaks in each set with median distances of 16.5 kb and 16.2 kb, respectively. We then scanned each set of peaks using the corresponding position weight matrices (PWMs) from HOCOMOCO v14 CORE^141^ or JASPAR 2026 CORE^142^ (vertebrates, non-redundant) and FIMO (MEME Suite 5.5.4), with a match significance threshold of p < 1×10^-4^. We scored each peak for the presence of the TF family if that peak contained at least one hit from any PWM belonging to the family; presence is binary, so multiple hits for the same family in one peak count once. We then compared the enrichment of each TF family between silencer-like and enhancer-like peaks with a two-sided Fisher’s exact test and calculated adjusted p-values using the Benjamini-Hochberg procedure.

### AlphaGenome, Borzoi, and Enformer predictions

We generated AlphaGenome^18^ predictions with the alphagenome-pytorch implementation (https://github.com/genomicsxai/alphagenome-pytorch), using the released all-folds checkpoint. We used the same 1 Mb context window centered on the *PPIF* TSS (hg38 GENCODE v46) as the reference allele for all predictions. We generated alternative alleles by modeling the real Swap-seq perturbation for each edit, deleting sequences between spacers and substituting them with the 71-bp barcode cassettes carrying corresponding barcodes. Because Swap-seq edits change sequence length and the model requires exactly 1,048,576 bp of input, we constructed alternative alleles by anchoring the 5’ edge of the context window and shifting the 3’ edge depending on the edit size — for example, a tile deleting 100 bp and inserting the 71-bp cassette shortens the locus by 29 bp, and to keep the input sequence length constant, we extended the 3’ end to include the 29-bp genomic DNA sequence downstream, and vice versa for edits that result in net insertion. We then scored the effects of Swap-seq edits on two output tracks. We took the THP-1 CAGE track on the *PPIF*-coding (+) strand and aggregated the predicted signal over a 100-bp window centered on the *PPIF* TSS, computing the log fold change of the signal of alternative allele over reference allele. Because Swap-seq edits shift the coordinates of the TSS, we realigned the output track of alternative alleles to reference coordinates and kept the CAGE quantification window constant. Similarly, we scored the edits with the CD14+ monocyte RNA-seq track, the closest available cell type to THP-1, using AlphaGenome’s gene-level scorer which aggregates predicted coverage over the exons of the *PPIF* gene, and calculated log fold change of alternative alleles over the reference allele. We averaged the barcode-level predictions to compare with the experimental results of each edit measured by Swap-seq.

To generate predictions for regulatory sequences in the lentiMPRA format, we assembled synthetic reporter cassettes that mirror the lentiMPRA construct architecture. We constructed two types of input sequences: (i) Promoter-to-CRE, each consisting of a promoter, the reporter gene, and the regulatory sequence; and (ii) LTR-to-LTR, all sequences between the 5’ and 3’ LTRs, including LTRs themselves. All sequences of promoters, regulatory elements, and plasmids were exactly as they were represented in the lentiMPRA (**table S4**). To define a baseline for each promoter, we constructed 30 negative-control sequences: 10 sequences designed to have no activities from a previous lentiMPRA library ^50^, and 10 independent dinucleotide-shuffles of each of the silencer s1 and the enhancer e2, which preserved the lengths and dinucleotide compositions of their sequences but destroyed TFBS. We predicted the activity of each sequence with the AlphaGenome THP-1 CAGE track with a fixed 4,096-bp or 8,192-bp context window for Promoter-to-CRE and LTR-to-LTR sequences, respectively. As our MPRA sequences are shorter than the context window, we embedded each MPRA sequence to the center of 20 background sequences, generated by dinucleotide-shuffled genomic sequences taken from neutral genomic regions. To quantify activities, we summed the predicted CAGE signals over the promoter plus 100 bp extending past the 3’ end. For each promoter, we defined the baseline as the average CAGE signals across the 30 negative controls, and computed each CRE’s effect as the log2 fold change of its CAGE over the negative control baseline, then averaged across the 20 backgrounds.

We generated predictions for Swap-seq edits using Borzoi^21^ and Enformer^19^ as described in Variant-EFFECTS^41^. Briefly, we used CD14+ monocyte CAGE track for both models, providing 524 kb and 196 kb input sequences centered at the *PPIF* TSS to Borzoi and Enformer, respectively. Two distal elements were excluded from the Enformer run as they were out of the 196 kb window. For both models, we quantified the model prediction as the log ratio of alternative to reference signal, aggregated over a 500-bp window centered on the *PPIF* TSS. We realigned the alternative allele output track to reference coordinates before aggregating signals, as for AlphaGenome.

### Artificial intelligence (AI) usage

Claude Opus 5 was used for writing and reviewing code for data analysis, and for editing the manuscript to improve clarity of language.

## Data Availability

The data generated in this publication, including *PPIF* Swap-seq screen, ATAC-seq, CUT&RUN, lentiMPRA, and silencer s1 Variant-EFFECTS screen, have been deposited in NCBI’s Gene Expression Omnibus and are accessible through GEO Series accession number GSE348496. All sequencing data associated with the THP-1 Micro-C and RCMC experiments with single clonal lines are available in the NCBI GEO under SuperSeries accession number GSE347552. All external datasets used for analysis are listed in **table S8**. The THP-1 ChromBPNet model used in this work is available at: https://zenodo.org/records/10403552.

## Code Availability

Swap-seq analysis pipeline is available on Github (https://github.com/EngreitzLab/Swap-seq) with all files used to process the current Swap-seq screen.

## Acknowledgements

We thank C. Ludwig for cloning pCL081 and pCL082, and Y. Finkel for cloning pYF023. We thank G. Martyn for helping with RNA FlowFISH and THP-1 cell culture. We thank B. Doughty for discussion about calculating Swap-seq effect sizes. We thank J. Linder for discussion about applying sequence-to-function models to study large deletions. We thank C. Rios-Martinez, G. Carattini, and A. Thurm for helping with flow cytometry and chemical inhibitor experiments. We thank P. Smaruj and G. Fudenberg for discussion about the Akita model. We thank J. Tan and B. Xia for discussion about the Chromnitron and C.Origami models. We thank D. Yao for feedback on designing dual-pegRNA constructs and choosing appropriate promoters. We thank N. Jain, Aradhana, J. Tycko, P. Du, and M. Bassik for helping with Swap-seq method development and screen designs. We thank D. Bredikhin for feedback on benchmarking AlphaGenome. We thank S. Conley for keeping the Engreitz Lab running and H. Bickford, M. Ameri, A. Parkkinen, and H. Emery for administrative support. We thank all the members of the Engreitz and Bintu Labs for discussions and feedback. We thank Stanford University and the Stanford Research Computing Center for providing computational resources and support as part of the Sherlock High-Performance Compute Cluster. We also thank the Stanford Genomics facility for providing Illumina MiSeq access and support.

## Author Contributions

X.S.C., M.T.M., and J.M.E. designed the study.

M.T.M. conceived of the Swap-seq method and optimized initial Swap-seq cassettes.

X.S.C., M.T.M., and J.M.E. designed the final Swap-seq constructs, cloning process, and conducted proof-of-concept experiments.

X.S.C., M.T.M., T.Z., and J.M.E. designed the Swap-seq *PPIF* screen.

X.S.C. and J.M.E. conducted Swap-seq *PPIF* screen and data analysis.

X.S.C., Z.C., A.K., and J.M.E. analyzed data with ChromBPNet.

M.N. conducted THP-1 Micro-C and RCMC. M.N. and X.S.C. analyzed the Micro-C and RCMC data.

A.S.H. supervised Micro-C and RCMC work.

X.S.C., A.N., Y.T., and J.M.E. designed, conducted, and analyzed the lentiMPRA experiment.

X.S.C., J.M.E., and L.B. designed and conducted the fluorescent reporter and chemical inhibition experiments, and analyzed flow cytometry data.

X.S.C., Z.G., and J.M.E. designed and conducted the silencer s1 Variant-EFFECTS experiment.

X.S.C., L.B., and J.M.E. conducted and analyzed ATAC-seq and CUT&RUN experiments.

K.O., X.S.C., A.K., and J.M.E. generated Borzoi and Enformer model predictions.

X.S.C., R.R., A.K., L.B., and J.M.E. conducted AlphaGenome benchmarking and analyzed the results.

M.U.S. and J.M.E. generated ENCODE-rE2G model predictions in THP-1 cells.

C.X. and S.C. generated single clonal cell lines and tested PRIME-Del with the supervision of X.S.C. and J.M.E.

J.M.E. and L.B. jointly supervised the work.

X.S.C., L.B., and J.M.E. wrote the manuscript with input from all authors.

