## Supplementary Information for "Systematic mapping of enhancers, silencers, and topological elements with prime editing deletion screens"

### Supplementary Figures

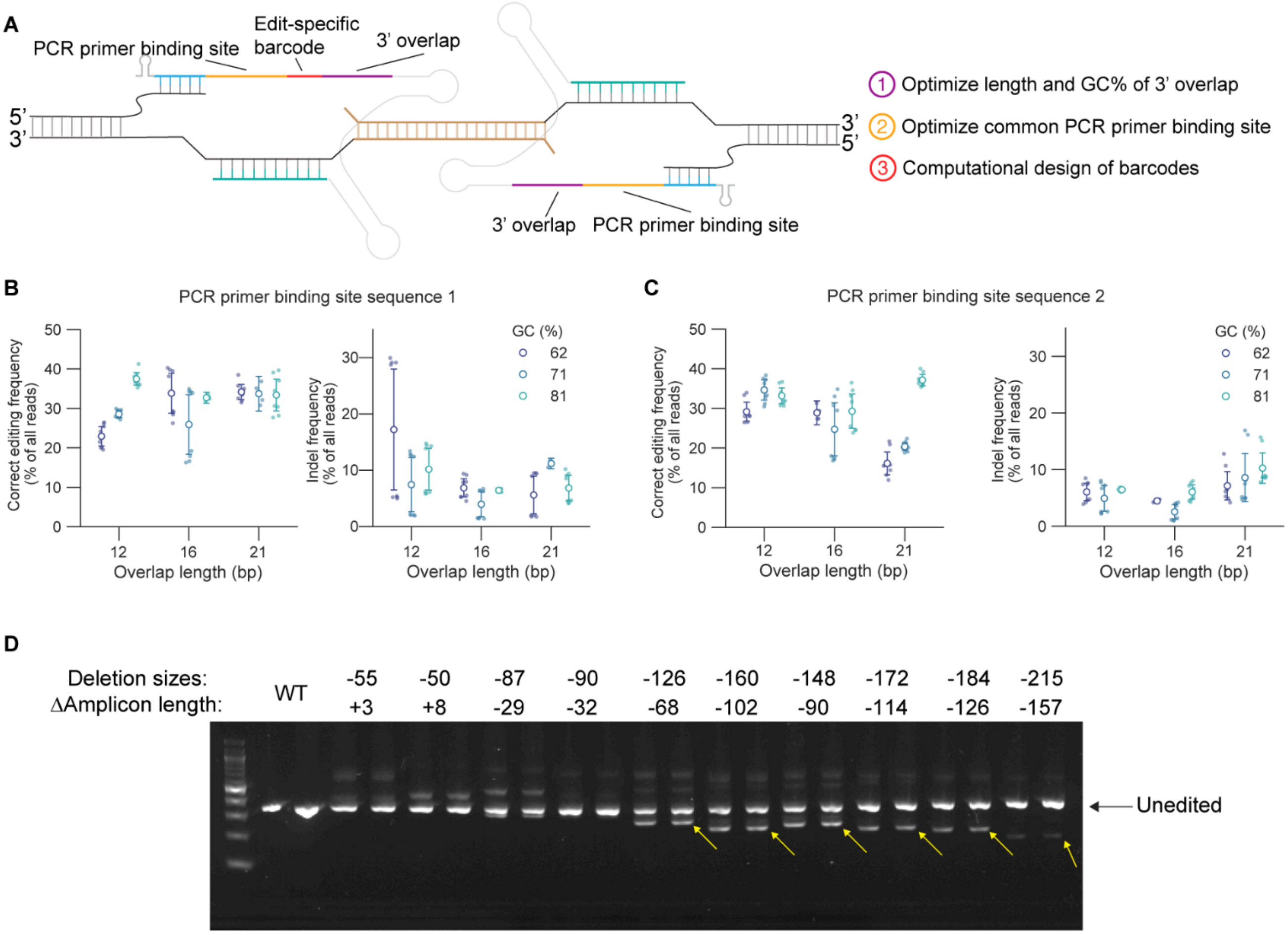


**Fig. S1. Optimization of the Swap-seq pegRNA RT templates**

**A.** Components on the Swap-seq pegRNA RT templates and optimization steps.

**B.** Testing length and GC% of 3' overlap on editing efficiency and indel rate, using sequence design 1. Aggregated between two loci, *HEK3* and *CCR5*. Error bars: 95% CI.

**C.** As in **B**, but with sequence design 2. Error bars: 95% CI.

**D.** Testing deletion sizes at an enhancer locus. PegRNAs encoding 55- to 215-bp deletions within an enhancer in the *PPIF* locus were transfected into HEK293T cells. Gel showing PCR amplification of the enhancer 72 h after transfection. Swap-seq deleted the target sequence and inserted a 58-bp barcode cassette, e.g., a 120-bp deletion will result in an amplicon 62 bp shorter than the WT.

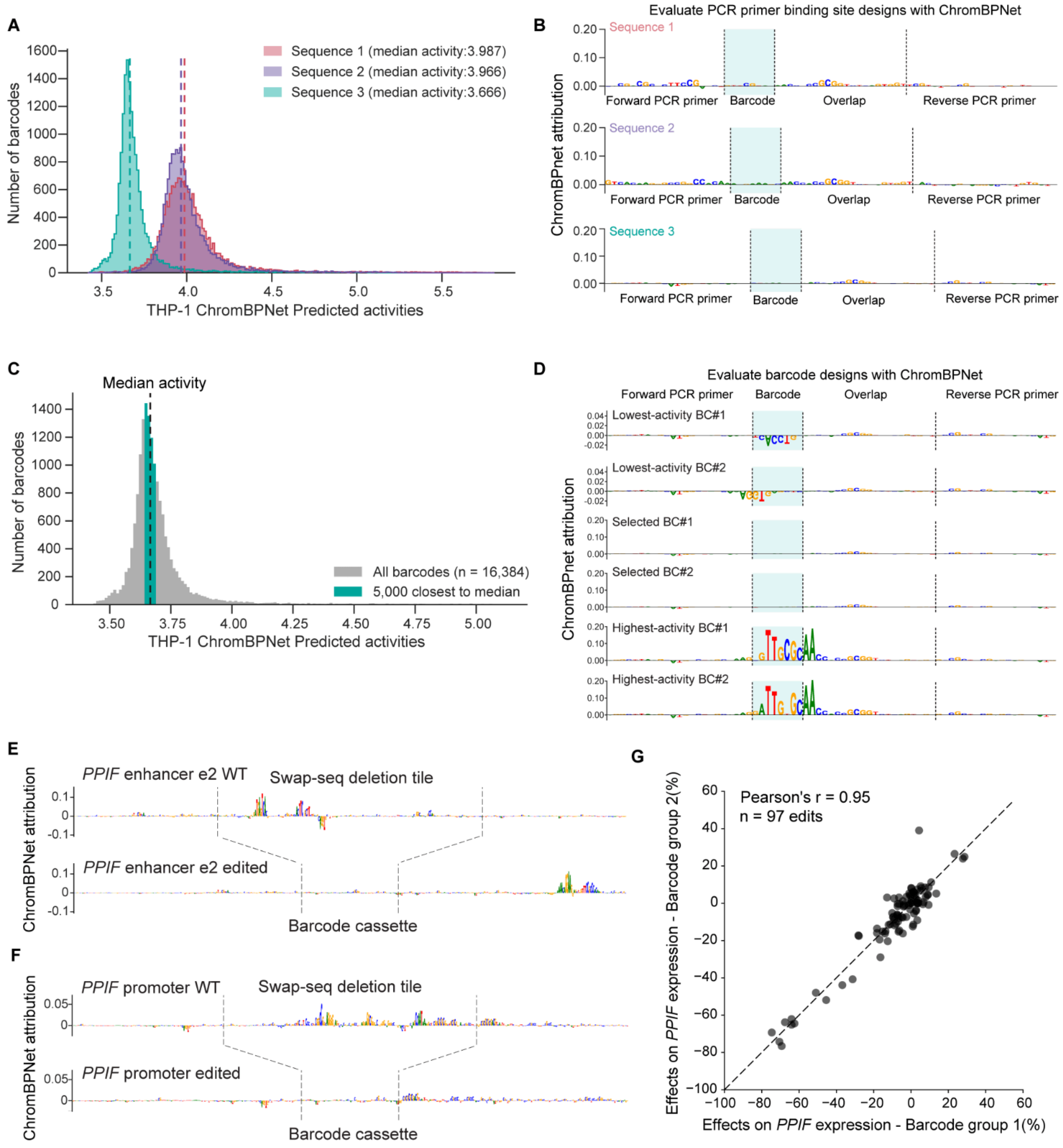


**Fig. S2. Computational designs of inactive Swap-seq barcode cassettes**

**A**. THP-1 ChromBPNet predicted activities of 3 PCR primer binding site designs with all possible 7-bp barcodes. Dashed lines: median activity of each PCR primer design. Activity scores are the log-transformed values of predicted ATAC-seq signals.

**B.** Evaluate primer binding site designs with ChromBPNet. THP-1 ChromBPNet attribution tracks were shown for three different PCR primer designs. For each design, the sequence with predicted activity closest to the median was chosen. The height of each letter represents the predicted contribution to accessibility of each base pair.

**C.** Evaluate barcode designs with ChromBPNet. 5,000 barcodes closest to the median activity were selected for designing pegRNAs. Dashed lines: median activity of barcode cassettes with PCR primer design 3 and all possible 7-bp barcodes.
**D.** ChromBPNet attribution tracks of exemplary barcode cassettes. Selected barcode cassettes are predicted to be inactive and devoid of TFBS (Middle: Selected BC#1 and 2). Excluded barcode cassettes have too low (Top: Lowest activity BC#1 and 2) or high (Bottom: Highest activity BC#1 and 2) predicted activities and contain TFBS.
**E.** *In silico* Swap-seq editing of an exemplary Swap-seq edit in enhancer e2. Top track: THP-1 ChromBPNet attribution track of the WT sequence centered at the deletion tile. Two dashed lines mark the left and right cut sites. Bottom track: After deletion, a Swap-seq barcode cassette will be inserted in place of the deleted sequence.
**F.** As in **E**, an additional example of *in silico* Swap-seq edit in the *PPIF* promoter.
**G.** Correlation of Swap-seq effects on *PPIF* expression split by barcode. From the screen, four barcodes of each edit were randomly split into two groups. Dots represent the average effects on *PPIF* expression from each barcode group. (Pearson's r=0.95, n = 97 edits with all four barcodes passing the frequency threshold of 0.001%).

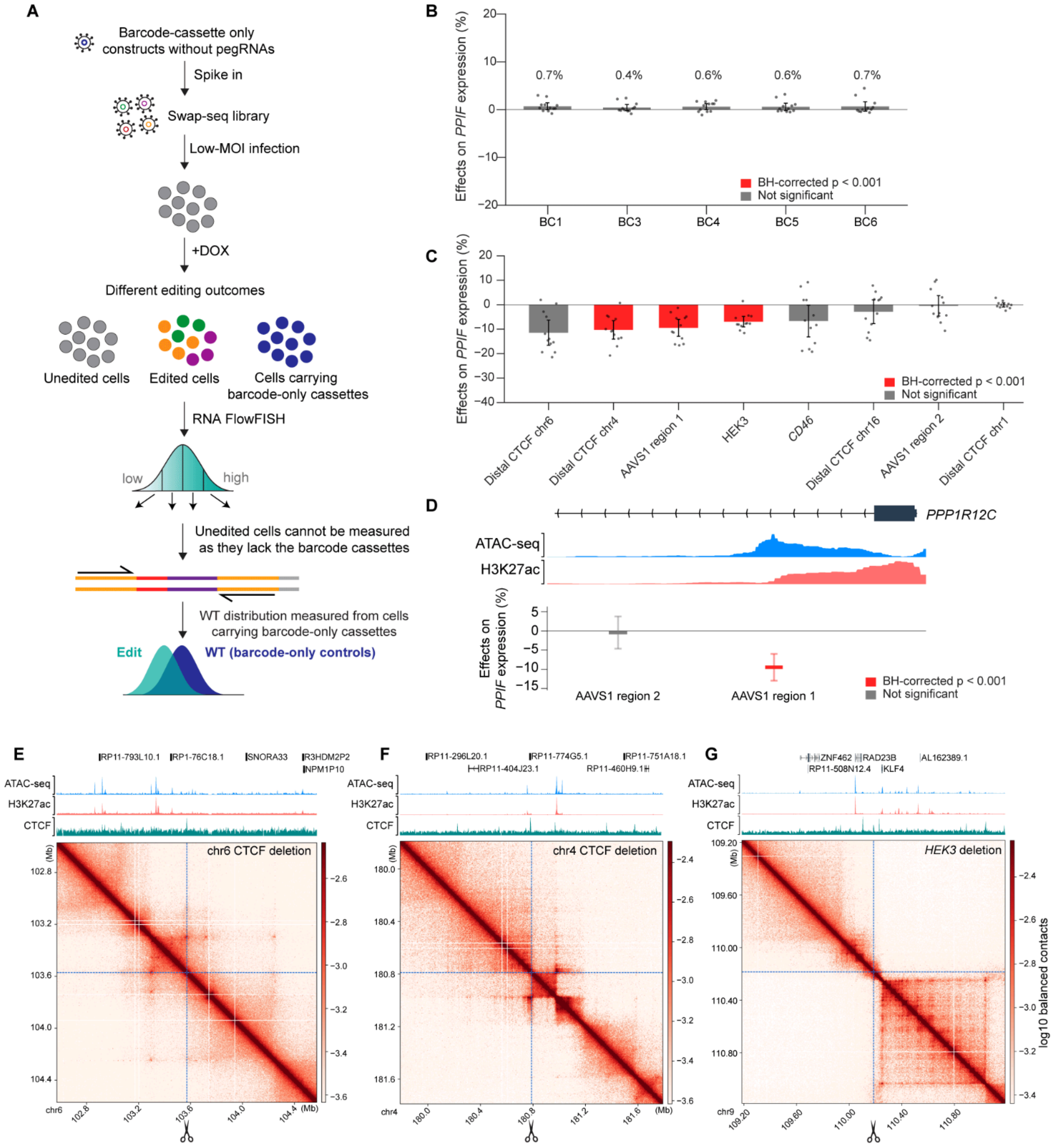


**Fig. S3. Negative controls for Swap-seq experiments**

**A**. Barcode-only constructs without pegRNAs were spiked into the Swap-seq library. As unedited cells do not contain a barcode cassette that can be sequenced, cells infected with barcode-only constructs were used as WT distribution to calculate the effects of each edit.

**B**. Negative control barcode-only cassettes have consistent close zero effects. Six barcode-only constructs, with different barcode sequences, were spiked into the Swap-seq library. Barcode #2 was arbitrarily chosen as the true “WT” and effect sizes were calculated for the remaining 5 barcode-only cassettes. Red, BH-corrected $p < 0.001$. Gray, not significant. Error bars: 95% CI.

**C**. Distribution of out-of-chromosome negative controls. Eleven edits targeting accessible regions outside of chromosome 10 were designed as additional negative controls. Eight edits passed the editing frequency threshold with their effect sizes shown. Red, BH-corrected $p < 0.001$. Gray, not significant. Error bars: 95% CI.
**D**. Swap-seq deletion effects at the *AAVS1* locus within the intron of *PPP1R12C*. Red, BH-corrected $p < 0.001$. Gray, not significant. Error bars: 95% CI.
**E-G**. Out-of-chromosome edits fall within CTCF binding sites and can alter 3D genome contact, leading to trans-effects on *PPIF*. Edits in (**E**) chromosome 6, (**F**) chromosome 4, and (**G**) *HEK3* locus overlap with CTCF binding sites that associate with TAD boundaries.

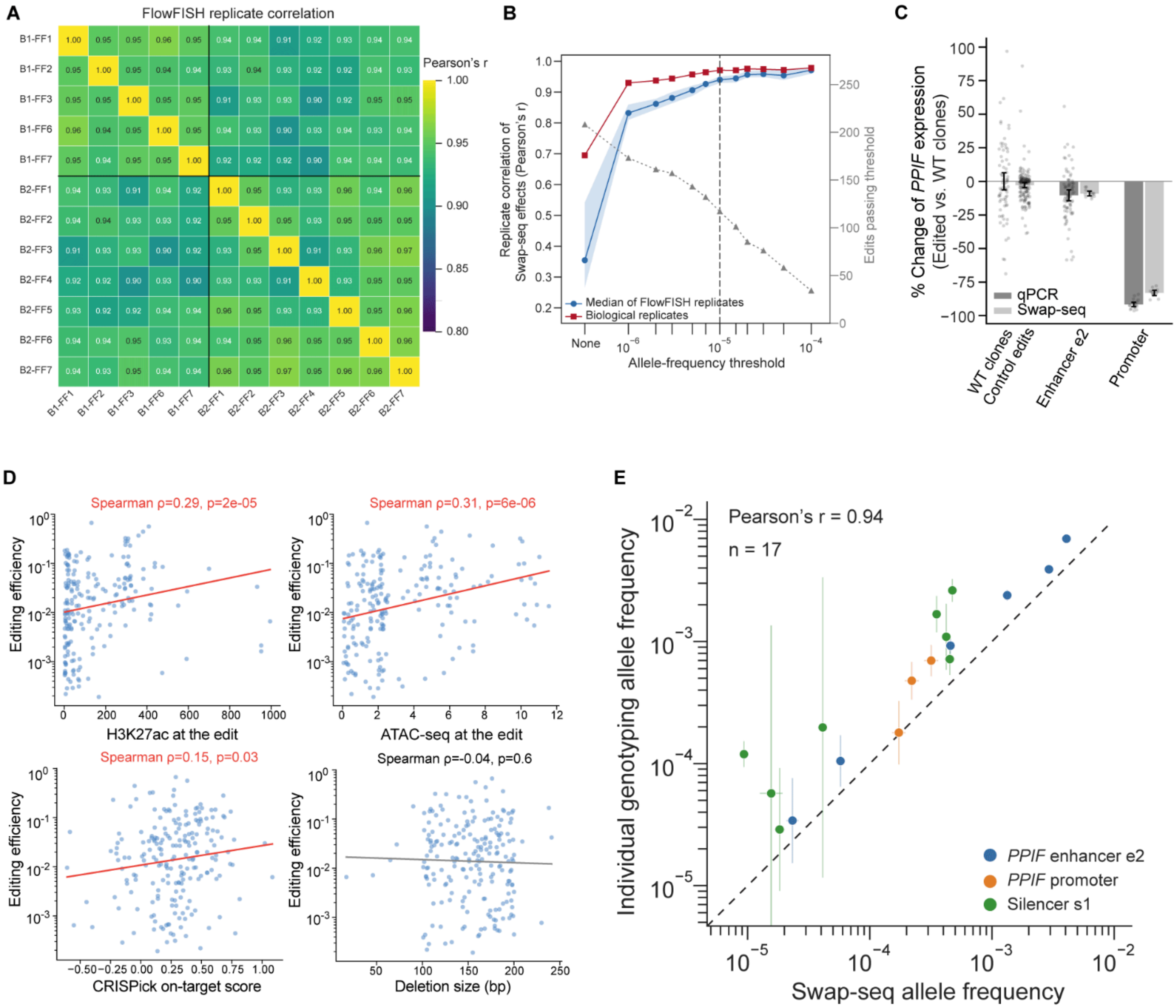

**Fig. S4. Technical validation of the Swap-seq screen at the *PPIF* locus**

**A**. Correlation of the effects of Swap-seq edits on *PPIF* expression between RNA FlowFISH replicates. FlowFISH replicates 4 and 5 in biological replicate 1 were excluded due to experimental errors.

**B**. Effects of allele frequency threshold on the correlation between biological and FlowFISH replicates. x-axis: allele frequency threshold; y-axis (left): Pearson's r between biological replicates or FlowFISH replicates; y-axis (right): number of edits passing threshold.

**C**. qPCR validation of Swap-seq effects. For qPCR, each dot represents a wild-type (WT) clone (n = 75) or a clone carrying homozygous edits in the enhancer e2 (n = 84) or promoter of *PPIF* (n = 12), averaged across 3 qPCR technical replicates. For Swap-seq, each dot represents an RNA FlowFISH replicate of negative control edits (n = 172), enhancer e2 edit (n = 12) or the *PPIF* promoter edit (n = 12). Swap-seq negative control edits include barcode-only controls and all out-of-chromosome edits. Error bars: 95% CI for the mean.

**D**. Correlation between editing efficiency and chromatin features and deletion size.

**E**. Correlation between direct genotyping and Swap-seq-estimated allele frequency at three loci, enhancer e2, *PPIF* promoter, and silencer s1. Error bars: 95% CI for the mean.

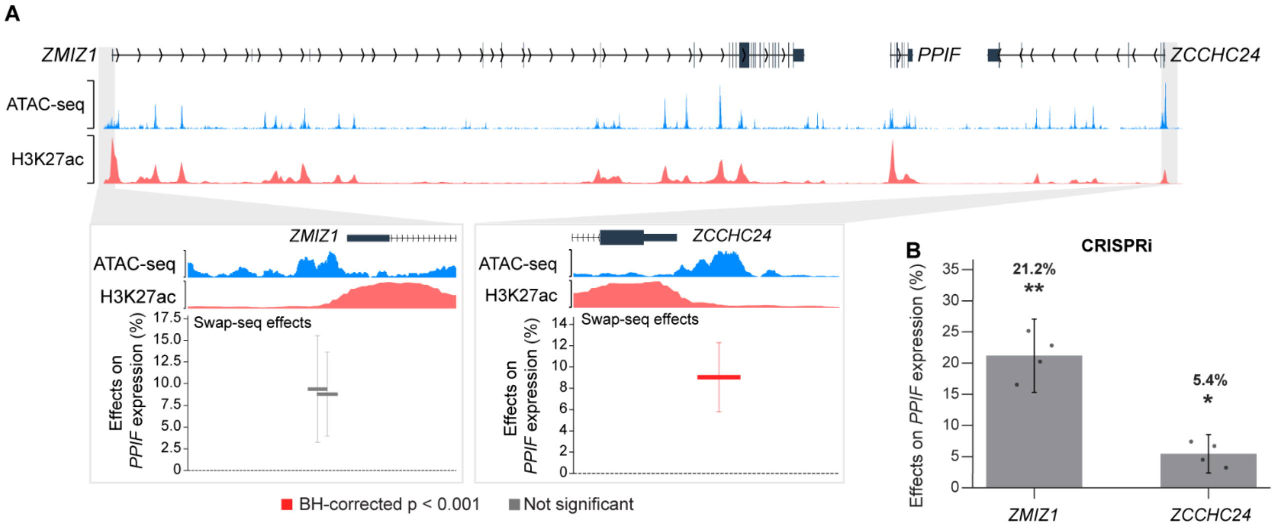


**Fig. S5. Perturbing the promoters of *ZMIZ1* or *ZCCHC24* increases *PPIF* expression**

**A**. Swap-seq effects of tiles within the promoter region of *ZMIZ1* and *ZCCHC24*. Red, BH-corrected p < 0.001. Gray, not significant. Error bars: 95% CI.

**B.** CRISPRi-FlowFISH effects of targeting the promoter of *ZMIZ1* and *ZCCHC24*. Each dot represents an RNA FlowFISH replicate. One-sample t-test: *p < 0.05, **p < 0.01. Error bars: 95% CI.

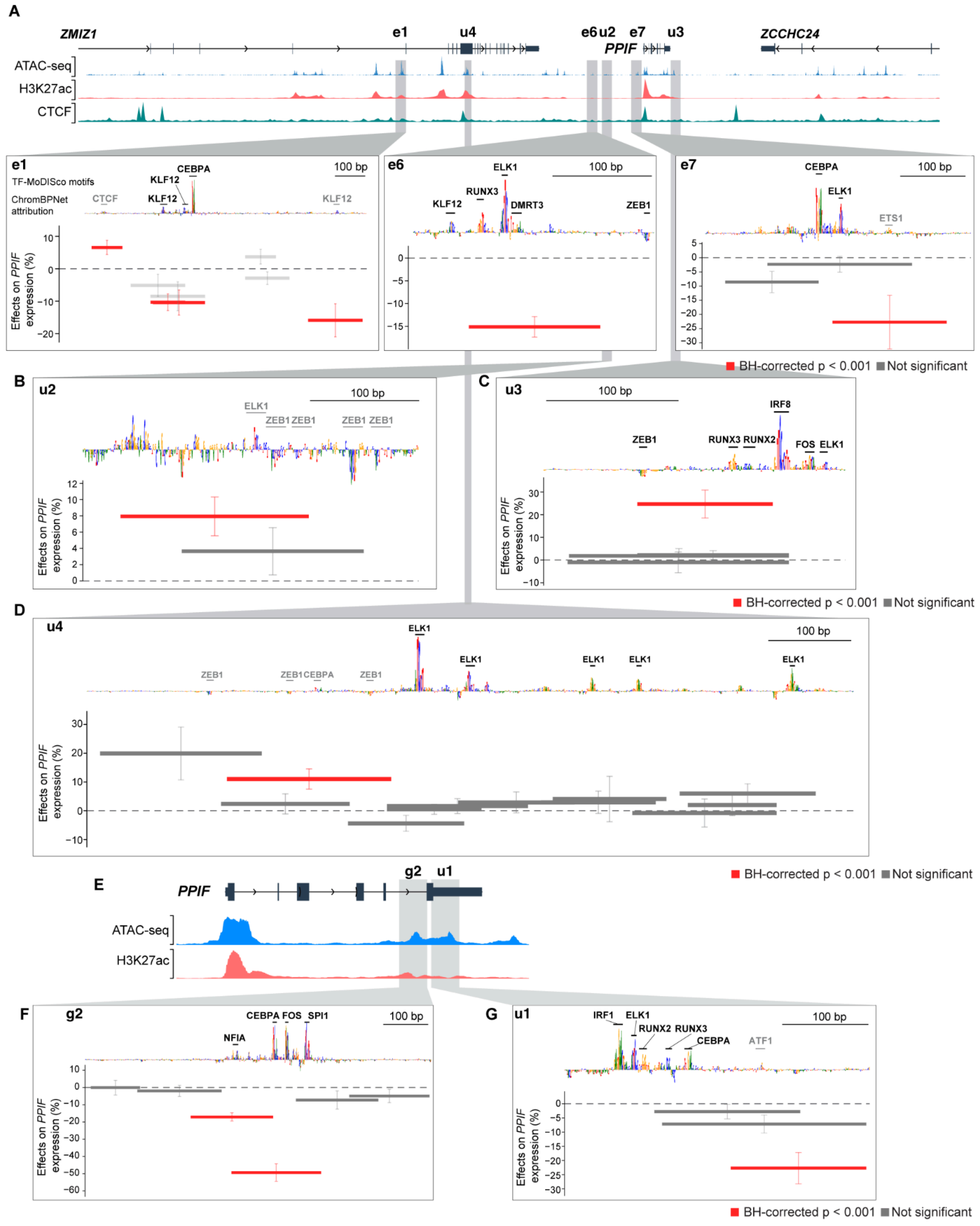


Fig. S6. Swap-seq discovers new regulatory elements

**A**. Swap-seq edits within three peaks (e1, e6, e7) led to a significant decrease of *PPIF*. Tiles with significant effects overlap with ChromBPNet predicted TFBS for activators. Red, BH-corrected $p < 0.001$. Gray, not significant. Error bars: 95% CI for the mean. Motifs in black are Fi-NeMo motifs with hit importance ≥ 0.1 (see **Methods**).

**B-D**. Swap-seq edits within three elements (u2-u4) led to significant increase of *PPIF* expression. (**B**) **u2**: Swap-seq tile with significant effects overlaps with predicted ZEB1 binding site. (**C**) **u3**: Swap-seq tile with significant effects overlaps with predicted ZEB1 and RUNX family binding sites. (**D**) **u4**: Swap-seq edit in a candidate element within the gene body of *ZMIZ1* led to upregulation of *PPIF*. The edit with significant effect overlaps with predicted ZEB1 and CEBPA binding sites. Red, BH-corrected $p < 0.001$. Gray, not significant. Error bars: 95% CI for the mean.

**E-G**. Two further edits within the *PPIF* gene body led to a significant decrease of *PPIF* expression. (**E**) Locus plot of the two edits in the *PPIF* gene body. (**F**) Swap-seq edit (g2) in an accessible element within intron 5 of *PPIF*. (**G**) Swap-seq edit (u1) in an accessible element within the 3’ UTR of *PPIF*. Red, BH-corrected $p < 0.001$. Gray, not significant. Error bars: 95% CI for the mean.

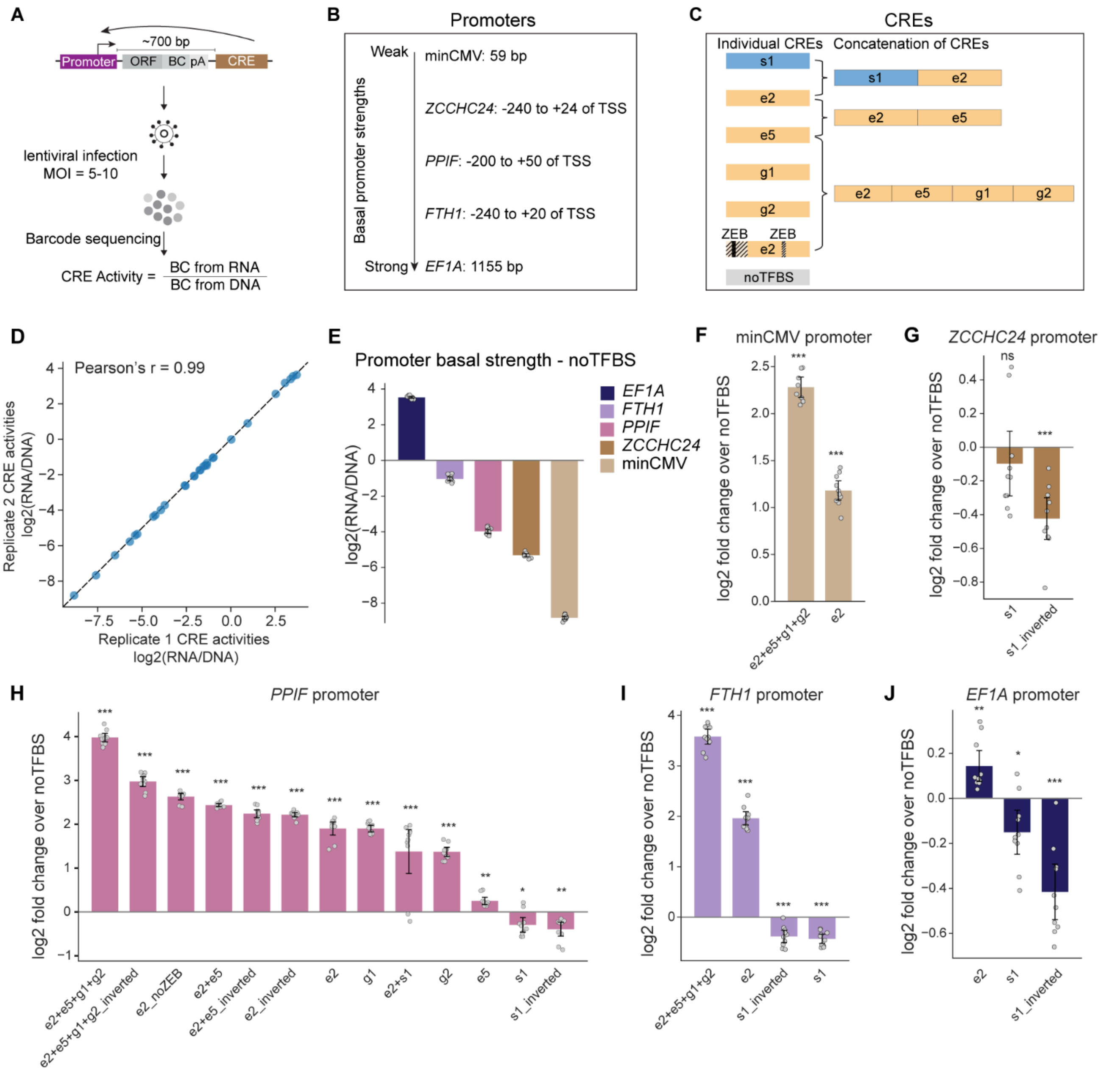


**Fig. S7. LentiMPRA measures regulatory effects of enhancers and silencer s1**

**A**. Schematic of the lentiMPRA workflow. Candidate CREs were cloned downstream of the promoter and transcripts to avoid transcriptional interference. We pooled individually made lentivirus, infected THP-1 cells at MOI of 5-10, and extracted DNA and RNA for barcode sequencing to determine the CRE activities.

**B**. Five promoters were selected to cover a range of basal promoter strengths.

**C**. The candidate CRE library contains both individual CREs and concatenations to test their interactions. For enhancer e2, we designed a version that removed the regions that contain potential repressor ZEB1 binding sites.

**D**. Correlation of CRE activities log2(RNA/DNA) of all promoter-CRE pairs between two biological replicates.

**E**. Activities of each promoter paired with the “noTFBS” negative control previously designed to be an inactive element. Each dot represents a corresponding barcode from each biological replicate. Error bars, 95% CI for the mean.
**F-J**. CRE activities tested on each promoter. All CRE activities were normalized to the activity of the “noTFBS” control paired with the corresponding promoter. (**F**) minCMV promoter, (**G**) *ZCCHC24* promoter, (**H**) *PPIF* promoter, (**I**) *FTH1* promoter, (**J**) *EF1A* promoter. Error bars, 95% CI for the mean. Two-sample t-test against the “noTFBS” negative control. *$p < 0.05$, **$p < 0.01$, ***$p < 0.001$, ns, not significant.

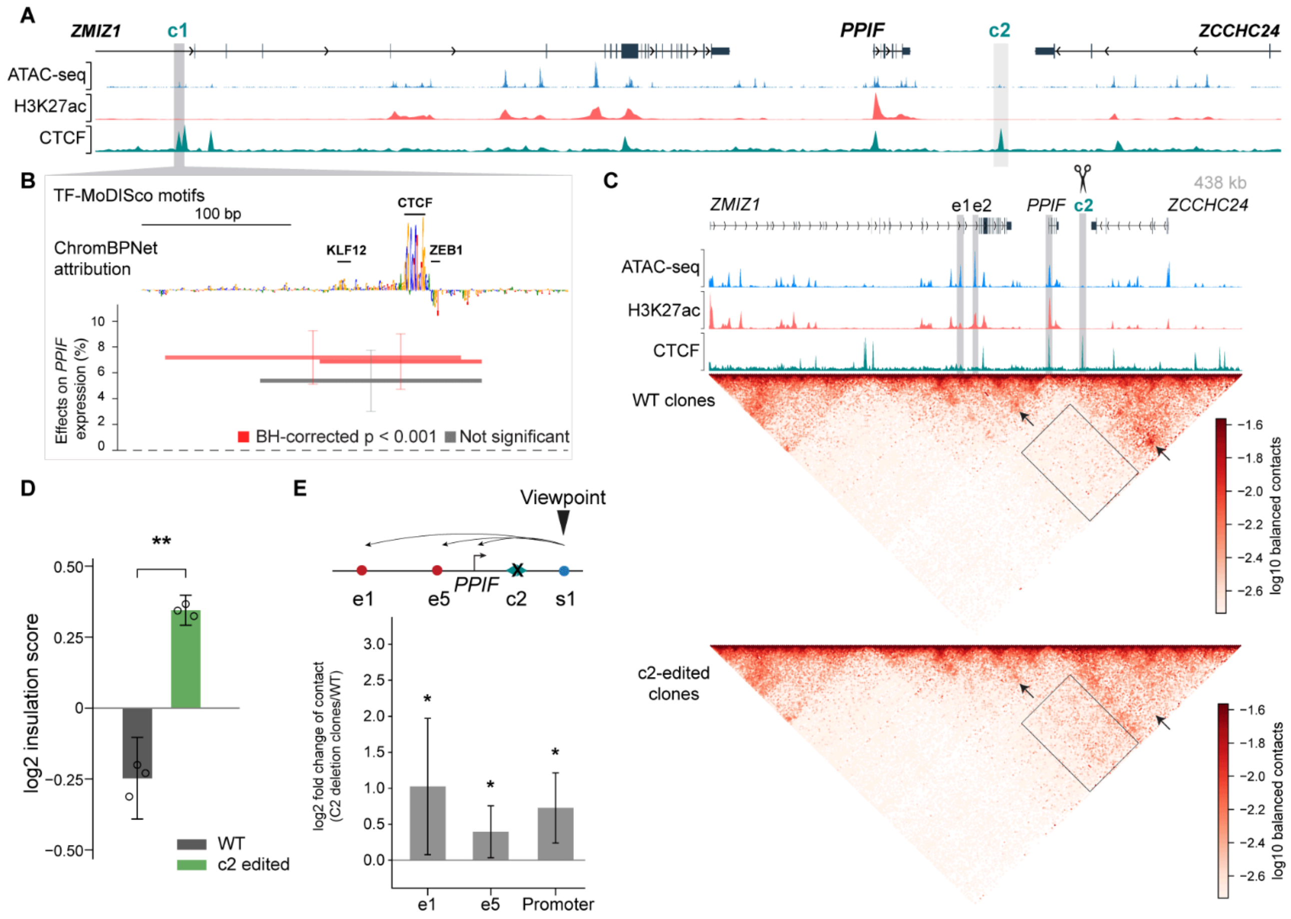


**Fig. S8. Swap-seq edits at CTCF binding sites altered the 3D genome structure**

**A**. Deletions within two CTCF binding sites (c1 and c2) led to significant changes in *PPIF* expression.

**B**. Swap-seq edits in c1. Red, BH-corrected $p < 0.001$. Gray, not significant. Error bars: 95% CI for the mean.

**C**. RCMC measurements of 3D genome structure at the *PPIF* locus (chr10:80,827,647–81,265,742) in WT (top) and edited clones (bottom). ICE-normalized contact maps shown at 1.6 kb binsize. Arrows highlight decreased enhancer contact with the *PPIF* promoter (left) or decreased CTCF-CTCF loop (right) after deleting c2. Box indicates the region with increased contact after deleting c2.

**D**. Insulation scores at c2, in WT and c2-deleted clones. Insulation scores were quantified using a 50 kb window. Each dot represents a WT clone or clones carrying homozygous c2 edit. Error bars: 95% CI for the mean. Welch's t-test: *$p < 0.05$, **$p < 0.01$.

**E**. Quantification of contact changes between WT and c2-deleted clones, with viewpoint on s1. Bars show log2 fold change of contact, averaged across three independent clones at 2 kb binsize. Error bars: 95% CI for the mean. Welch's t-test: *$p<0.05$, **$p<0.01$.

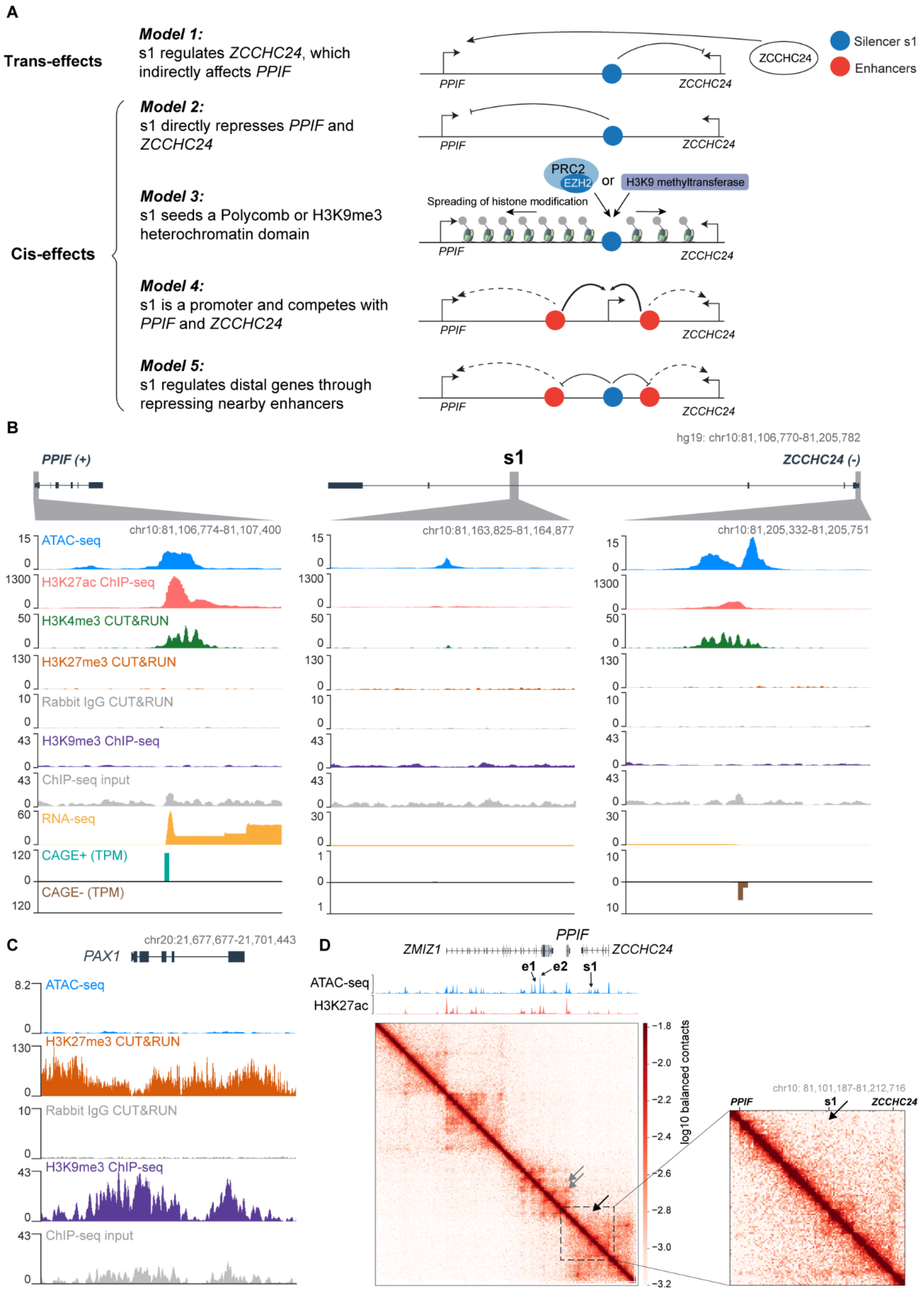
A
Trans-effects
Model 1:
s1 regulates ZCCHC24, which indirectly affects PPIF
Cis-effects
Model 2:
s1 directly represses PPIF and ZCCHC24
Model 3:
s1 seeds a Polycomb or H3K9me3 heterochromatin domain
Model 4:
s1 is a promoter and competes with PPIF and ZCCHC24
Model 5:
s1 regulates distal genes through repressing nearby enhancers
Silencer s1
Enhancers
PPIF
ZCCHC24
PRC2
EZH2
or
H3K9 methyltransferase
Spreading of histone modification
B
hg19: chr10:81,106,770-81,205,782
PPIF (+)
s1
ZCCHC24 (-)
chr10:81,106,774-81,107,400
chr10:81,163,825-81,164,877
chr10:81,205,332-81,205,751
ATAC-seq
H3K27ac ChIP-seq
H3K4me3 CUT&RUN
H3K27me3 CUT&RUN
Rabbit IgG CUT&RUN
H3K9me3 ChIP-seq
ChIP-seq input
RNA-seq
CAGE+ (TPM)
CAGE- (TPM)
C
chr20:21,677,677-21,701,443
PAX1
ATAC-seq
H3K27me3 CUT&RUN
Rabbit IgG CUT&RUN
H3K9me3 ChIP-seq
ChIP-seq input
D
PPIF
ZMIZ1
ZCCHC24
e1
e2
s1
ATAC-seq
H3K27ac
log10 balanced contacts
chr10: 81,101,187-81,212,716
PPIF
s1
ZCCHC24

**Fig. S9. Models of silencer s1 mechanism and genomic analysis**

**A**. Potential models of the silencer s1 repression mechanisms. Model 1: s1 deletion affects *PPIF* through trans-effects. Deleting s1 led to an increase of *ZCCHC24*, which indirectly affected *PPIF* expression. Model 2: s1 directly interacts with and represses the promoters of *PPIF*. Model 3: s1 interacts with protein complexes such as PRC2 and/or methyltransferases that deposit H3K27me3 and/or H3K9me3 locally, and the spreading of histone modifications can lead to repression of distal genes. Model 4: s1 is a promoter that competes with *PPIF* and *ZCCHC24*. Model 5: s1 represses nearby enhancers to regulate distal genes.

**B**. Experimental data show limited H3K27me3, H3K4me3, H3K9me3, or transcription activities at s1. (left) *PPIF* promoter, (middle) silencer s1, (right) *ZCCHC24* promoter.

**C**. Exemplary locus with high H3K9me3 and H3K27me3 signals.

**D**. THP-1 Micro-C contact map at the silencer s1 locus at 2 kb resolution after balanced normalization. Gray arrows: contact between the *PPIF* promoter and enhancers e1 and e2. Black arrow: s1 does not have focal contacts with the promoter of *PPIF*. Zoomed-in map showing Micro-C contact map in the range of chr10: 81,101,187-81,212,716 (hg19) at 1 kb resolution.

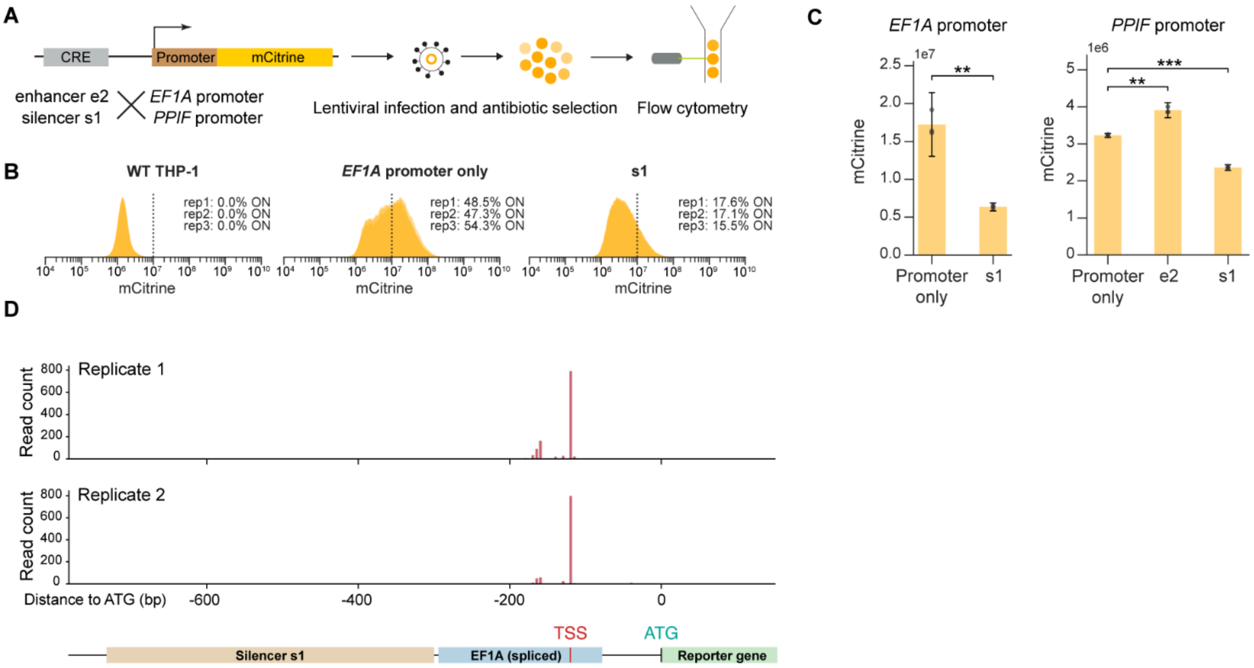


**Fig. S10. 5' TSS mapping with fluorescent reporter**

**A**. Set up of the fluorescent reporter assay to test the repressive effect of the silencer. Silencer s1, or a positive control enhancer e2, were cloned upstream of promoters and delivered into cells through lentiviral integration. Gene expression was quantified by measuring cell fluorescence level with flow cytometry.

**B**. Distribution of fluorescence signals of (1) WT THP-1 cells, (2) THP-1 cells infected with *EF1A*-only constructs, and (3) cells infected with a construct with s1 cloned in front of the *EF1A* promoter.

**C**. Silencer s1 led to a significant decrease of fluorescence signal of both *EF1A* and *PPIF* promoters. Error bars: 95% CI for the mean. Welch's t-test: $^{**}p<0.01$, $^{***}p<0.001$.

**D**. 5' TSS mapping of the fluorescent construct showed expected TSS and no transcription initiation from the silencer s1. Each bar represents the number of aligned reads that initiate at the corresponding location in the reporter constructs.

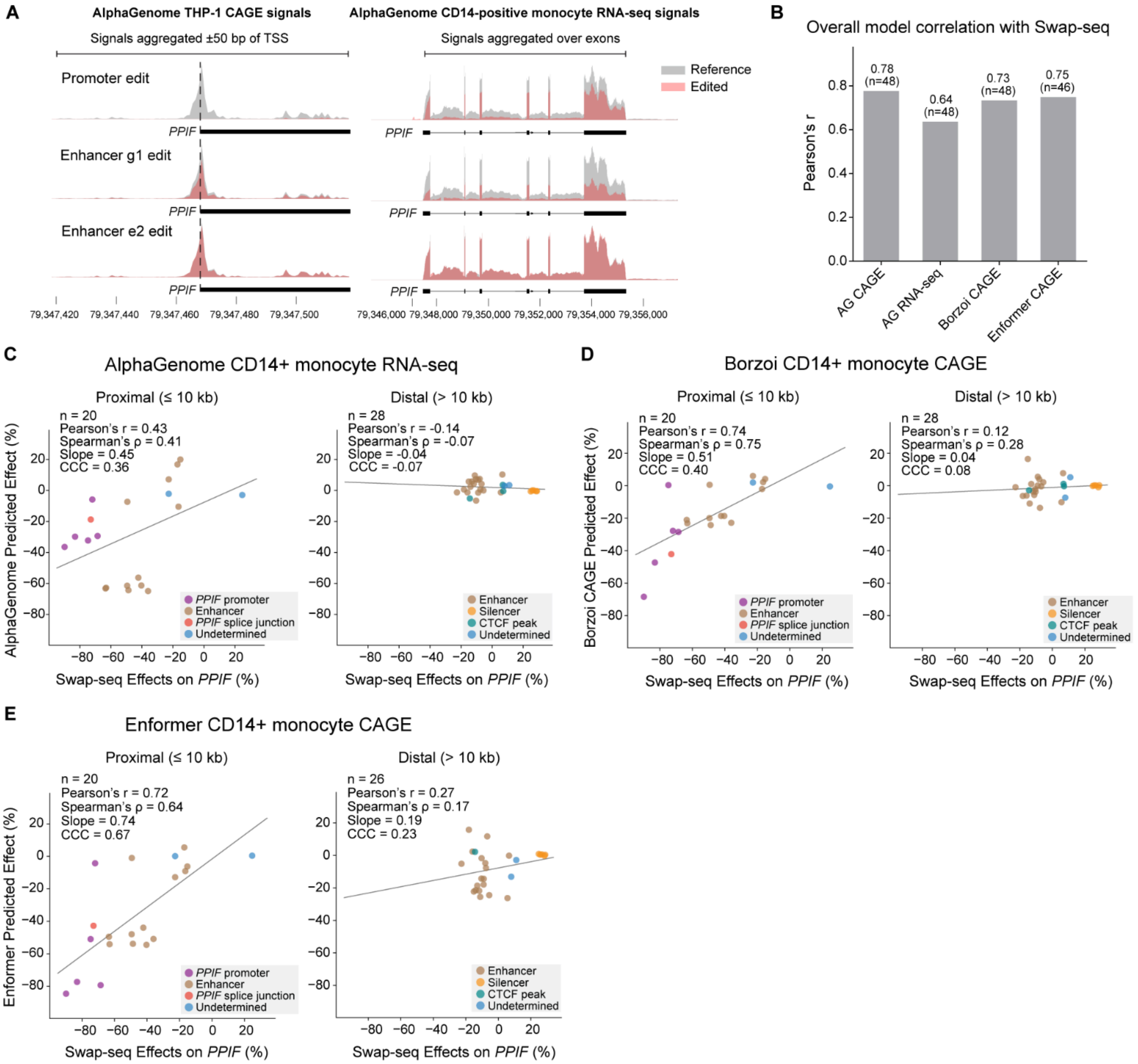


**Fig. S11. Benchmarking sequence-to-function models with the Swap-seq data**

**A**. AlphaGenome predicted CAGE (left) and RNA-seq (right) signals for three Swap-seq edits in the *PPIF* promoter, g1, and e2. Gray: reference sequence; Red: sequence containing the corresponding Swap-seq edit. Coordinates on the x-axis are in hg38.

**B**. Overall Pearson's correlation coefficient between Swap-seq effects and model predictions.

**C-E**. Correlation between Swap-seq effects and model predictions, stratified by distances of edits to the TSS. Left: Distance to TSS ≤ 10 kb; Right: Distance > 10 kb. (**C**) AlphaGenome CD14+ monocyte RNA-seq track (1-Mb context). (**D**) Borzoi CD14+ monocyte CAGE track (524 kb). (**E**) Enformer CD14+ monocyte CAGE track (196 kb).

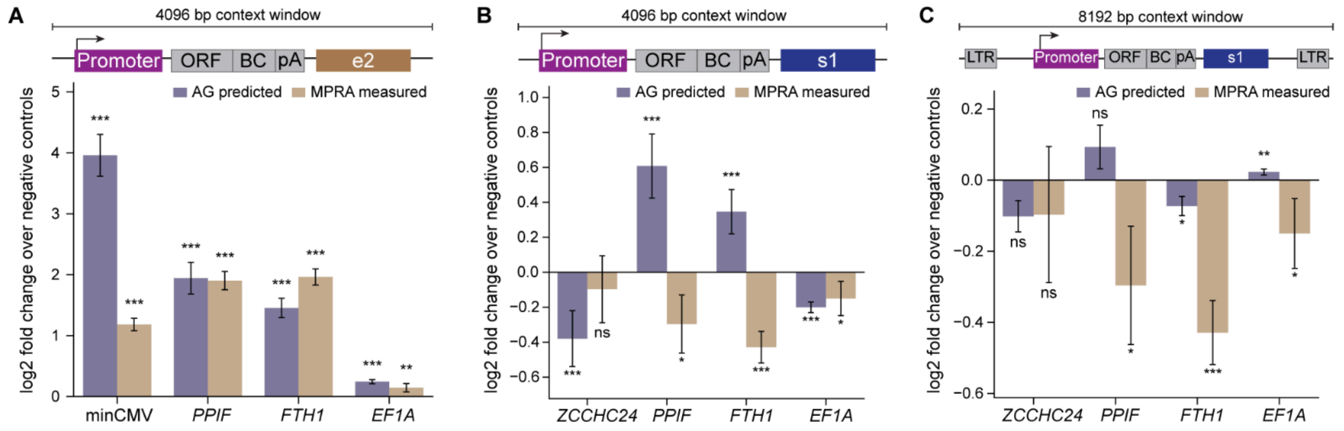


**Fig. S12. Comparing AlphaGenome predictions with lentiMPRA**

**A.** Comparing AlphaGenome THP-1 CAGE predictions with lentiMPRA measurements of enhancer e2 on promoters. Schematic on top shows the configuration of lentiMPRA sequences used as input for prediction and the size of the AlphaGenome context window. Each MPRA sequence was embedded in 20 different background sequences to reach the target context size. Bars represent log2 fold change over negative controls, averaged across the 20 sequences. Error bars: 95% CI for the mean. Two-sample Welch's t-test: *p<0.05, **p<0.01, ***p<0.001, ns, not significant.

**B**. As in (**A**), comparing AlphaGenome THP-1 CAGE predictions with lentiMPRA measurements of silencer s1 on promoters. Error bars: 95% CI for the mean. Two-sample Welch's t-test: *p<0.05, **p<0.01, ***p<0.001, ns, not significant.

**C.** As in (**B**), but including the lentiMPRA sequence from 5' to 3' long terminal repeats (LTRs) as the input sequence for AlphaGenome prediction. Error bars: 95% CI for the mean. Two-sample Welch's t-test: *p<0.05, **p<0.01, ***p<0.001, ns, not significant.

### Supplementary Tables

**Table S1. Individual pegRNAs and oligo nucleotides used in this work**
**Table S2. Swap-seq *PPIF* tiling screen pegRNA library sequences**
**Table S3. Swap-seq *PPIF* tiling screen results**
**Table S4. MPRA sequences and data**
**Table S5. Silencer s1 Variant-EFFECTS pegRNAs and data**
**Table S6. Silencer-like peaks containing at least three of four motifs**
**Table S7. Model prediction of Swap-seq edits on *PPIF* expression**
**Table S8. External datasets**
